# Characterising AlphaFold 3’s ability to predict T cell antigen specificity

**DOI:** 10.64898/2026.07.08.737208

**Authors:** Benjamin McMaster, Ali El Moselhy, Ilija Ilievski, Christopher J. Thorpe, Nicole L. La Gruta, Jamie Rossjohn, Charlotte M. Deane, Hashem Koohy

**Affiliations:** Medical Research Council Translational Immune Discovery Unit (MRC TIDU), Weatherall Institute of Molecular Medicine, University of Oxford, United Kingdom; Department of Statistics, University of Oxford, United Kingdom; Open Targets, Wellcome Genome Campus, United Kingdom; European Molecular Biology Laboratory, European Bioinformatics Institute (EMBL-EBI), United Kingdom; Infection and Immunity Program and Department of Biochemistry and Molecular Biology, Monash University, Australia; Institute of Infection and Immunity, Cardiff University, United Kingdom

## Abstract

T cells are a key part of the adaptive immune system. Using their surface-bound T cell antigen receptors (TCRs), these cells scan peptides and other antigens presented to them by major histocompatibility complex molecules (MHCs) on the surface of cells, searching for abnormalities. Although determining the map between TCRs and their target antigens is of vital importance for the design of safe and effective T cell-based vaccines and therapeutics, decoding these interactions is challenging. Experimental methods are not scalable, and sequence-based computational methods have issues generalising to new antigens. The IMMREP25 benchmark of methods for predicting T cell antigen specificity showed that AlphaFold-based methods promise improved generalisation to novel antigens. However, the ability of structure prediction models to predict T cell antigen specificity has not been robustly evaluated previously. In this work, we characterise AlphaFold’s ability to predict T cell antigen specificity. We created a pipeline for high-throughput prediction of TCR:peptide-MHC (pMHC) structures using AlphaFold that is *>* 100 fold faster than the default implementation and used it to benchmark AlphaFold 3 (AF3) and similar models at predicting T cell antigen specificity. We investigated the underlying correlates of AlphaFold-derived binding scores and found that the model’s predictive power is related to the positioning of TCRs over the pMHC and not chemical interactions. Furthermore, we refine the AlphaFold-derived binding scores by training a machine learning model we call the PAE Aggregator. We then investigate AF3’s ability to uncover the clustering rules of TCR repertoires and recapitulate mutational scanning experiments. These analyses show that AlphaFold3 clusters sequence-similar TCRs according to their binding mode and detects disrupting point mutations accurately. Our results highlight both the promise and the current limitations of structure-based approaches for predicting TCR specificity, guiding the development of more reliable immunological prediction methods.

## Introduction

T cells play a key role in our adaptive immune system. These cells use their surface-bound T cell antigen receptors (TCRs) to scan peptides and other antigens presented to them by major histocompatibility complex molecules (MHCs) and related homologues, enabling them to identify threats to the body. T cells take on many roles within the immune system: killing infected cells, recruiting other cell types in the fight, or dampening overly active immune responses. No matter the cell function, antigen recognition is central to the role of a T cell. It is desirable to understand this recognition to design better vaccines against infectious diseases, create new therapeutics for cancers, and uncover the mechanisms underlying autoimmune disorders. However, understanding the antigen specificity of T cells is challenging. The stochastic development of TCRs through V(D)J recombination [1] leads to *≈* 10^18^ unique TCRs across the human population [2]. There can be *≈* 10^12^ possible peptides presented to these cells, and the MHC region of the human genome is the most diverse locus with over 40,000 alleles [3] and a distinct set of 12 in each person. This challenge is exacerbated by the many-to-many relationship TCRs have with peptide-MHCs (pMHCs) through cross-reactivity and common specificity [4]; one TCR can bind multiple pMHCs, and similarly, one pMHC can be bound by multiple unique TCRs.

The combinatorial complexity of this problem makes current experimental methods intractable, and thus, efforts in the last decade have turned to computational approaches due to their ability to uncover complex patterns in complex feature spaces [5]. Many machine learning models to predict T cell antigen specificity have been developed using amino acid sequence data [6, 7, 8, 9]; however, independent benchmarks have shown that these models fail to generalise beyond the restricted set of epitopes with interacting TCR information present in public databases [10, 11, 12]. This problem has been dubbed the “unseen epitope challenge” and means that these sequence-based models provide little help in deorphanising TCRs that arise in disease-specific studies. As a result of this and some promising work from Bradley [13], we speculated that exploring alternative data modalities and modelling approaches, such as protein structures and AlphaFold, respectively, may allow for generalised predictions of T cell antigen specificity [14]. The IMMREP25 benchmark, specifically aimed at addressing the unseen epitope challenge, demonstrated the potential of AlphaFold-based approaches [15]. Almost all models achieving better than random performance in this benchmark employed AlphaFold 3 (AF3) [16] or derivative models [17, 18, 19].

While AlphaFold predicts the 3D structure of a protein from its amino acid sequence (or other biomolecular formats), it also generates confidence scores to accompany its predictions. These include predicted local-distance difference test (pLDDT), a per-atom (or residue in the case of AlphaFold 2 (AF2) [20]) confidence score for estimating how accurate the local placement of the atom is, and predicted aligned error (PAE), a residue-to-residue confidence score estimating the global accuracy of residue placements relative to one another. Other confidence scores, such as the predicted template modeling (pTM), interface predicted template modeling (ipTM), and ranking scores, are derived from these model outputs to determine aggregate properties of the predictions. In their work, Bradley [13] shows that these confidence scores can discriminate between binding and non-binding TCR:pMHCs, as the binders are scored more confidently than the non-binders. This predictive ability powered Bradley [13] to the top of the IMMREP25 benchmark [15].

Based on the success of AF2 and AF3, other models have been developed following similar principles for biomolecular structure prediction. Boltz (1 and 2) [18, 19] and Chai [17] are general structure prediction models, and TCRBuilder (2 and 2+) [21, 22] and TCRmodel2 [23] are TCR and TCR:pMHC specific structure predictors derived from AlphaFold [14]. While AF2 has a permissive software licence, AF3 imposes more restrictions for commercial use, and these other models aim to offer an alternative. Independent benchmarks of these models have been conducted for TCR:pMHC structure prediction accuracy [24, 25], but little has been done to evaluate their ability to predict antigen specificity.

In this work, we aim to systematically characterise AF3 and like models’ ability to predict T cell antigen specificity. We build a platform to enable high-throughput predictions of TCR:pMHC structures from sequences using AF3. We then benchmark AF2, AF3, Chai-1, and Boltz-2 at predicting T cell antigen specificity and TCR:pMHC complex structures, and determine that AF3 has the best performance on these tasks. Next, we determine the important input signals AF3 uses for predicting T cell antigen specificity and structure predictions. Having established AF3 as a generalised predictor of antigen specificity, we investigate the correlates of its ability to predict antigen specificity with features from crystallography data and show that its predictive ability is strongly associated with the relative placement of the TCR over the pMHC and not chemical interactions between them. Next, we enhance AF3’s predictive ability of T cell antigen specificity by designing a simple model that learns the importance of equivalent TCR residue-pMHC residue pairs and removes effects associated with intrinsic TCR and pMHC confidence that we call “PAE Aggregator”. We then investigate AF3’s ability to uncover the rules of antigen-specific TCR clustering using our derived structural features. Finally, we use our PAE Aggregator score to detect disrupting amino acid mutations from experimental data of CD4+ T cells binding coeliac antigens. Our work highlights AF3’s ability to predict T cell antigen specificity and address the unseen epitope challenge as well as other immunological challenges, while also demonstrating its limitations on these tasks.

## Results

### Evaluating structure prediction as a framework for predicting T cell antigen specificity

#### Accelerating MSA and template generation

Using default settings, running an AF3 prediction can take approximately an hour for a single TCR:pMHC complex, making large volumes of predictions intractable. To overcome this problem, we developed an accelerated pipeline (described in “Accelerated data pipeline for TCR:pMHC structure prediction”) that reduces prediction time from hours to minutes. The critical bottleneck lies in MSA generation, where the default AF3 settings require an average of around 51 minutes per complex for this stage alone (see Figure 1A). Our pipeline, built on TCRmodel2, reduces MSA generation to an average time of 29 seconds, yielding a 100-fold speedup, shifting the computational bottleneck away from MSA building to the inference stage. Using the TCRmodel2 sequence databases also reduces the storage required from around 300 GB to 50 MB. We also explored using the MMSeqs2 [26] MSA generation pipeline employed by ColabFold [27]; however, we found that it was not optimised for our hardware and led to longer MSA generation times, on average around 3 hours per MSA.

**Figure 1:**
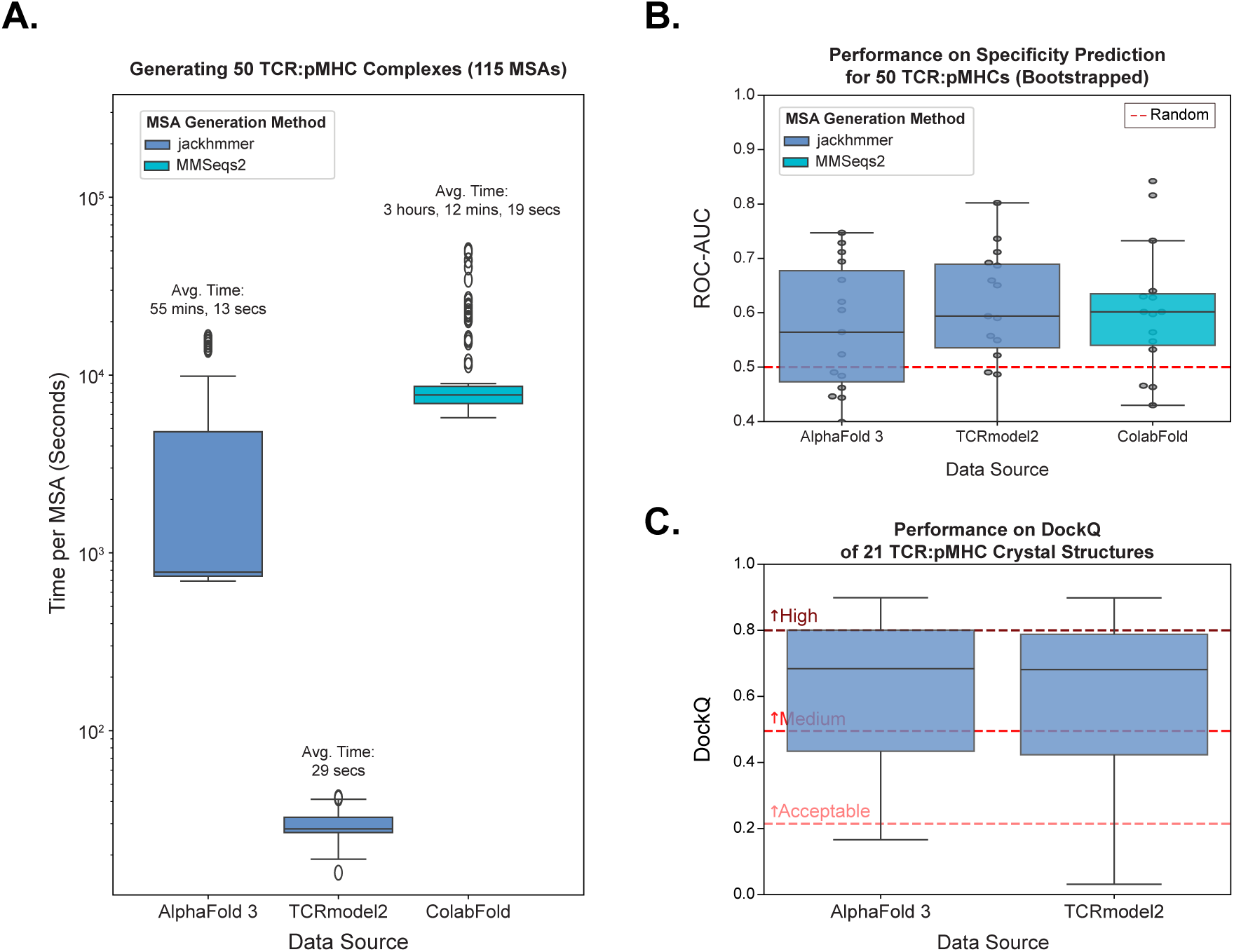
Effect of multiple sequence alignment (MSA) strategy choice on performance. **A.** Generation time to generate the necessary MSAs for 50 TCR:pMHC complexes. **B.** Boostrapped Area under the receiver operating characteristic curve (ROC-AUC) performance predicting the specificity of 50 TCR:pMHC interactions. Random performance is annotated by the red dashed line. **C.** DockQ performance of predicting 21 AF3-holdout TCR:pMHC structures. The critical assessment of predicted interactions (CAPRI) grading of DockQ scores is annotated.

On a small set of 50 TCR:pMHC sequences (25 binders and 25 non-binders), we investigated whether using the accelerated MSA generation pipeline compromised AF3’s ability to predict T cell antigen specificity. Highlighted in Figure 1B (and Figure S1A), we demonstrate that using our accelerated MSA generation pipeline achieves the same performance as the standard AF3 MSA generation pipeline (non-significant p-values for an analysis of variance (ANOVA) test). We then tested whether our accelerated MSA pipeline affected the quality of predicted structures, the main task AlphaFold was trained to perform. We tested the different approaches on our holdout set of 21 TCR:pMHC structural complexes (see “Collecting structure data”). Again, as Figure 1C demonstrates (and Figure S1B), we see no statistically significant difference between our MSA pipeline and the AF3 pipeline (non-significant p-values for a Wilcoxon signed-rank test).

Using the accelerated MSA generation pipeline gave a > 100 fold speed-up compared to the out-of-the-box AF3 MSA generation pipeline with no trade-offs in either antigen specificity prediction or structure prediction tasks. These modifications allow for a TCR:pMHC complex to be predicted in a few minutes and allow for more in-depth analysis comparing the predictive ability of these models as well as structure prediction-based analysis of medium-sized repertoires of TCRs and pMHCs.

#### Benchmark of AlphaFold 3 against open-license structure prediction models used in IMMREP25

We aimed to compare AF3 with open-licensed models AF2, Boltz, and Chai that were used in the IMMREP25 benchmark to predict the specificity of T cell antigens. Here, we used an expanded set of 1500 TCR:pMHC complex sequences (750 binders and 750 non-binders) [15]. We used the same MSAs and templates for all models (except AlphaFold2, which has limited support for paired MSA representations) that were generated by our accelerated MSA pipeline. Figure 2A (and Figure S2A) shows that Chai-1 and AF3 have comparable ability to predict T cell antigen specificity, and both of these models outperform the predecessor AlphaFold2. Boltz-2 only had comparable predictive ability to AF2, despite being based on the AF3 architecture. The differences in performance were assessed using an ANOVA test at a significance level of 0.05. These tests yielded a p-value of 1.90 *×* 10^−19^. Post hoc statistical testing was done using independent t-tests with an adjusted significance threshold using a Bonferroni correction.

**Figure 2:**
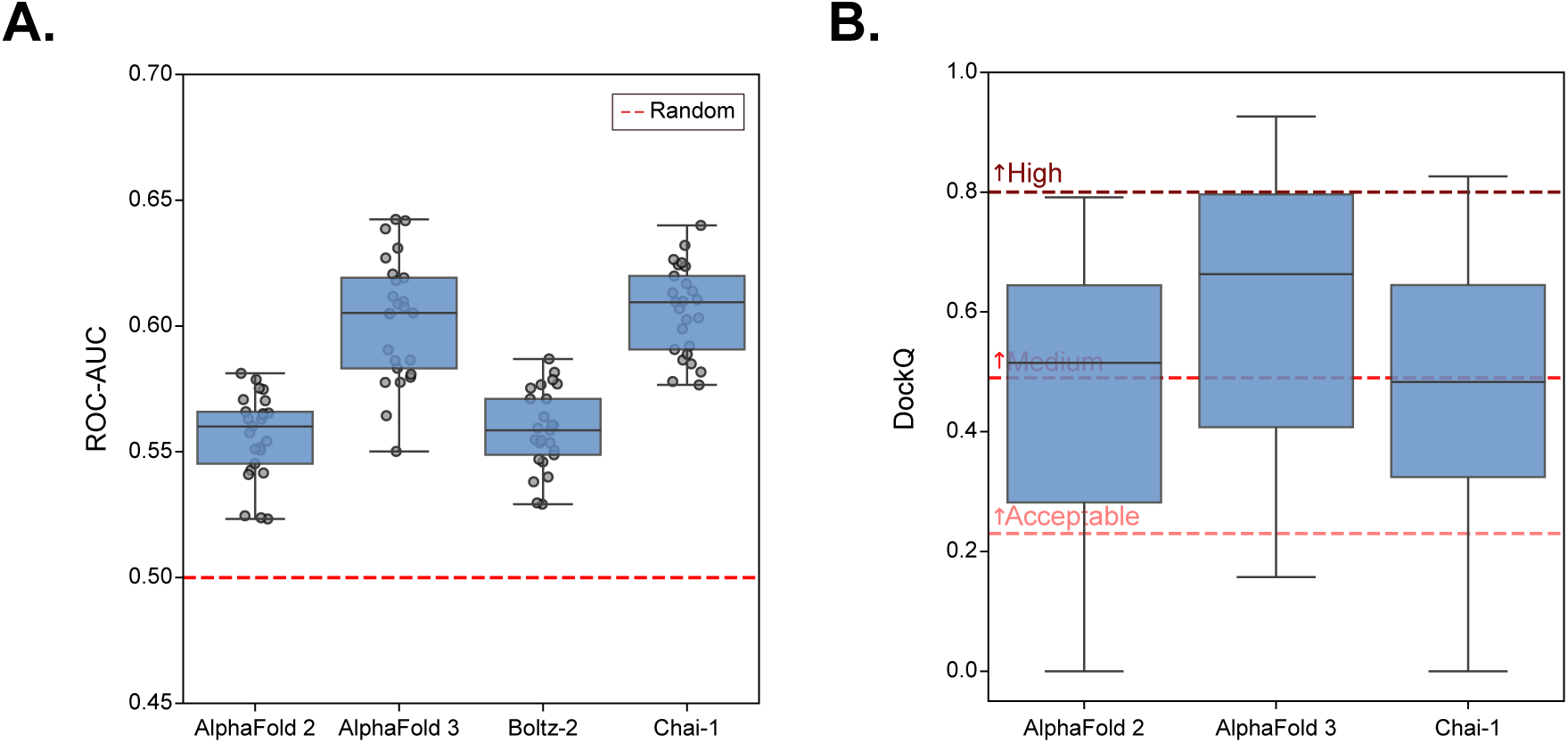
Benchmark of structure prediction models. **A.** ROC-AUC performance at predicting antigen specificity from 1500 TCR:pMHCs. **B.** DockQ performance at predicting TCR:pMHC complex structures from benchmark set. Boltz-2 was not included as only three structures were within the model’s training cut-off. The CAPRI grading of DockQ scores is annotated.

Turning to the structure prediction benchmark, we see that AF3 achieves the best performance on the holdout set of 21 TCR:pMHC structures (see Figure 2B and Figure S2B). A Kruskal-Wallis test shows significant differences (p-value of 2.27 *×* 10^−5^ at a significance level of 0.05), and post hoc testing showed that AF3 had a significant increase compared with AF2 and Chai-1 (assessed by Wilcoxon rank-sum tests and adjusted p-values using the Bonferroni correction). Qualitatively, when looking at the breakdown of CAPRI classifications in Figure 2B, we can see that AF3 has the highest portion of predicted structures designated as “High” quality. Of note, Boltz-2 was not included as only three structures (protein data bank (PDB) IDs 8shi, 8i5c, and 8i5d) were not in the training data for the model. Performance of at predicting these structures is shown in Figure S3.

Through this benchmarking, it is clear that AF3 has made improvements over AF2 at predicting T cell antigen specificity as well as the reported improvements in structure prediction. Alternative models, such as Chai-1, can achieve near-comparable results but fall short on some tasks.

#### Ablations of AlphaFold 3’s input channels

Along with sequence information, AF3 provides three additional signals to the machine learning model to generate 3D protein complex structures: unpaired MSAs, paired MSAs, and structure templates [16]. For each chain in the protein complex, the unpaired MSAs provide co-evolutionary information about which residue mutations co-occur within that chain. The paired MSAs then provide information about which residue mutations co-occur across different chains of the protein complex. The templates provide scaffolds of similar proteins for each protein chain individually. We sought to understand the impact each of these input signals provides for predicting T cell antigen specificity and the overall structural complex.

To determine the contributions from each of these input signals, we compared the performance on our dataset of 1500 TCR:pMHC interactions and 21 benchmark structures for various ablations of the input channels. Figure 3 (and Figure S4) depicts the results of these ablations. The ablations show that MSA signals are essential for both T cell antigen specificity prediction and structure prediction. Conducting an ANOVA test shows significant differences across all ablations in predicting T cell antigen specificity (p-value of 1.54 *×* 10^−69^ at a significance level of 0.05), and a Kruskal-Wallis test showed significant differences in predicting the structure quality (p-value of 2.02 *×* 10^−85^). Through post hoc analysis of all pairwise combinations using two-tailed independent t-tests with adjusted p-values using the Bonferroni correction, we see that excluding both the unpaired MSAs and the paired MSAs leads to a significant reduction in performance in predicting T cell antigen specificity across all comparisons. Additionally, the addition of templates does have a minor impact, as the same post hoc analysis shows there is a significant increase observed between only including templates and not including any of the three additional input signals. A non-parametric version of the same post hoc analysis was done using Wilcoxon rank sum tests for the structure quality evaluation, which also reached the same conclusions. In the case where all three input signals are removed, the sequence information alone is not enough for the model. The performance at predicting T cell antigen specificity drops to random (ROC-AUC of *≈* 0.5) and the structure prediction quality degrades completely (DockQ of *≈* 0.0). The similar trends observed for the T cell antigen specificity evaluation and structure prediction benchmark indicate the connected nature of predicting T cell antigen specificity and high-quality structure predictions, also observed by Deleuran and Nielsen [28].

**Figure 3:**
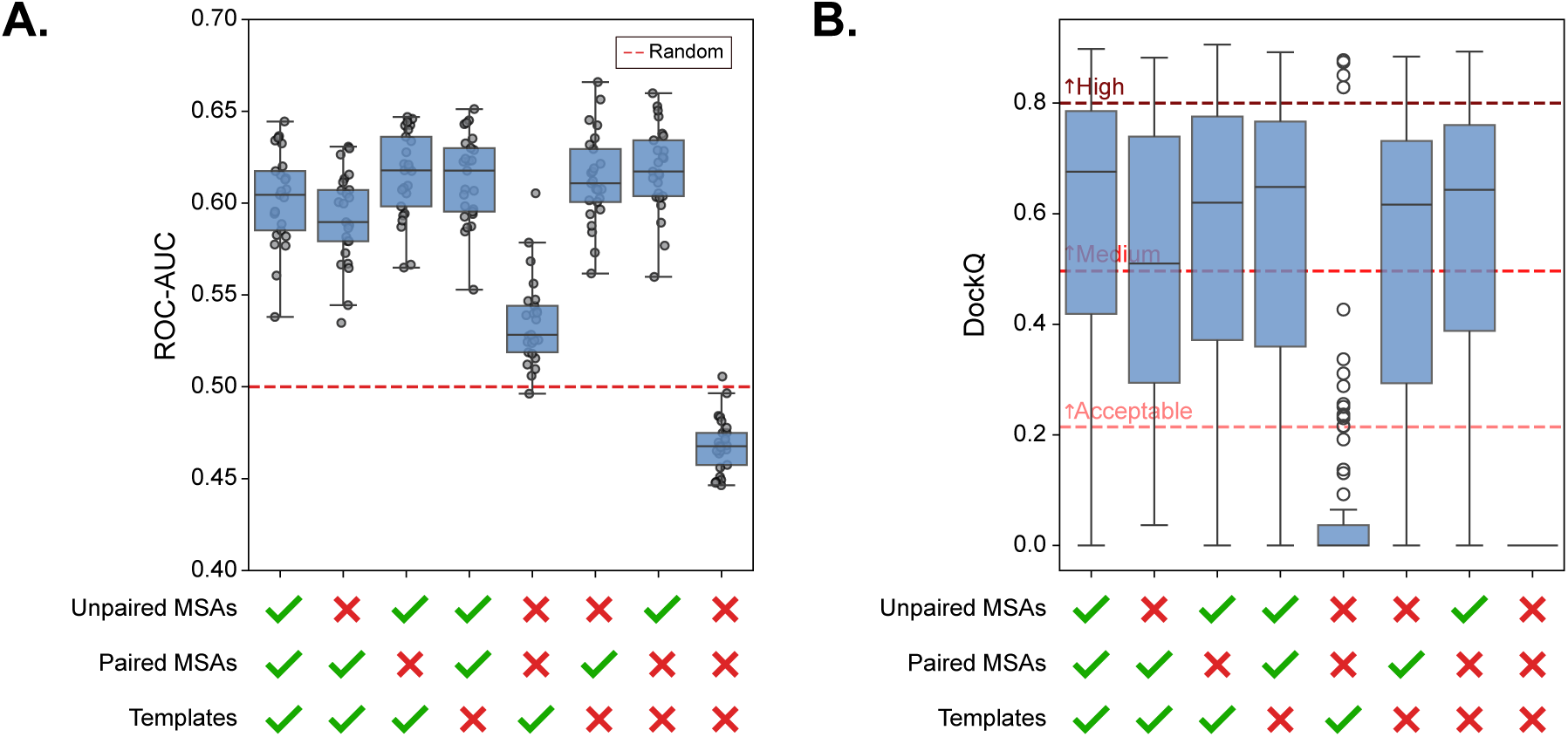
Effects of AF3 input channel ablations on performance. **A.** ROC-AUC performance at predicting T cell antigen specificity using 1500 TCR:pMHC interactions. **B.** DockQ performance at predicting TCR:pMHC complex structures. The CAPRI grading of DockQ scores is annotated.

From this analysis, we establish the importance of the MSA information for AF3’s ability to predict T cell antigen specificity and give evidence that the template information also plays a minor role. For that reason, we include all three signals in our further analysis using AlphaFold to predict T cell antigen specificity. The minor differences between including paired and unpaired MSA information potentially indicate that these signals are not fully optimised for this task, as the binding interface between TCRs and pMHCs is only partially evolutionarily driven, and exact specificity comes from stochastic pairings between the complementarity determining region (CDR) loops and peptides.

### Structural correlates of AlphaFold 3-derived binding scores

Having established AF3 as a powerful generalised predictor of T cell antigen specificity, we aimed to understand the underlying drivers of its performance. Described in “Parameterising TCR:pMHC complexes”, we parameterised our structural dataset of 251 experimental TCR:pMHC complexes with the following attributes following a similar process to Quast et al. [30]: pitch angle, tilt angle, roll angle, scanning angle, X distance, Y distance, Z distance, shape complementarity, buried surface area, number of interactions, number of interaction types, and contact maps (Figure 4A). From the distributions of these different features of TCR:pMHCs, we fit appropriate statistical distributions to score TCR:pMHC complex structures on measures of canonical binding. The parameters for these distributions are listed in Table 1. Not included in this table, the probability densities of the contact maps were computed using the normalised proportion of contacts in TCR:class I peptide-MHC (pMHC-I) and TCR:class II peptide-MHC (pMHC-II) complexes separately. For all residue contacts, the total proportion of contacting residues in the 251 structures was computed and then renormalised between class I and class II MHC complexes. To calculate the probability of an individual TCR:pMHC complex’s contacts, we take the mean of the reported normalised proportions of interacting residue pairs.

**Figure 4:**
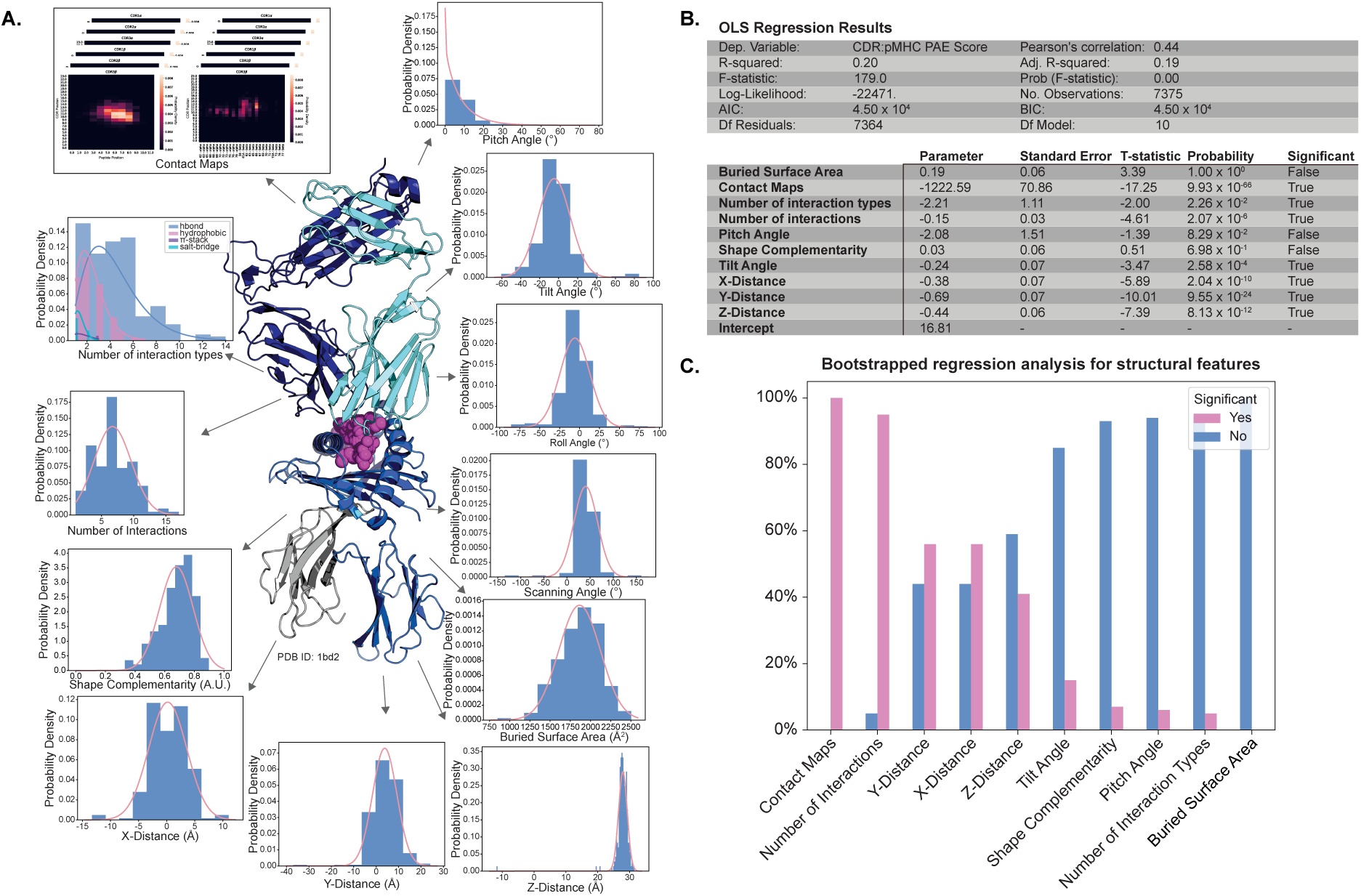
Correlation of TCR:pMHC structural features to AlphaFold confidence scores. A. Structural features of TCR:pMHC interactions and distributions derived from structures selected from STCRDab [29]. **B.** Results of a multiple linear regression between the selected structural features and the CDR:pMHC PAE Score. **C.** Bootstrap analysis of the structural feature correlates to the CDR:pMHC PAE Score.

**Table 1:** Parameterisation of TCR:pMHC structural features from 251 crystallography structures selected from the STCRDab.

| Feature | Distribution Type | Parameters |
| --- | --- | --- |
| Scanning Angle | Normal | $\mu = 40.47, \sigma = 25.61$ |
| Pitch Angle | Gamma | $\alpha = 0.93, x = 0, \theta = 8.96$ |
| Tilt Angle | Normal | $\mu = -4.70, \sigma = 17.17$ |
| Roll Angle | Normal | $\mu = -5.89, \sigma = 18.72$ |
| X-Distance | Normal | $\mu = 0.18, \sigma = 3.40$ |
| Y-Distance | Normal | $\mu = 3.82, \sigma = 5.47$ |
| Z-Distance | Gaussian Mixture | $\vec{w} = \begin{pmatrix} 0.004 \\ 0.996 \end{pmatrix}, \vec{\mu} = \begin{pmatrix} -11.79 \\ 27.90 \end{pmatrix}, \vec{\sigma} = \begin{pmatrix} 0.001 \\ 1.37 \end{pmatrix}$ |
| Shape Complementarity | Normal | $\mu = 0.68, \sigma = 0.11$ |
| Buried Surface Area | Normal | $\mu = 1864.63, \sigma = 258.05$ |
| Number of Interactions | Normal | $\mu = 6.71, \sigma = 2.90$ |
| Number of Interaction Types: |  |  |
| Hydrogen bond | Gamma | $\alpha = 3.27, x = 0, \theta = 1.33$ |
| Hydrophobic | Gamma | $\alpha = 3.94, x = 0, \theta = 0.60$ |
| π-stack | Gamma | $\alpha = 3.57, x = 0, \theta = 0.48$ |
| Salt-bridge | Gamma | $\alpha = 8.17, x = 0, \theta = 0.16$ |

We then scored our dataset of 1500 TCR:pMHC complexes predicted with AF3 (7500 total complexes) using these distributions derived from crystallography data. To understand the correlates of AF3’s confidence score, we fit a multiple linear regression model between the CDR:pMHC PAE Score and the structural feature probabilities (described in further detail in “Selecting structural features and fitting multiple linear regression model” and summarised in Figure 4B). The linear model achieved a Pearson’s correlation of 0.44 between the selected structural features and the CDR:pMHC PAE Score. To assess the structural correlates of the AF3-derived confidence scores, we analysed the individual contributions of each feature to the overall linear model in a bootstrapped analysis. Described in “Bootstrap anaylsis of structural correlates to AF3 confidence scores” and depicted in Figure 4C, our bootstrap analysis showed that the contact maps, general number of interactions, X distance, and Y distance of the TCR over the pMHC were highly correlated to the AF3 confidence scores (features achieving significance in > 50% of bootstrap trials). The interaction types, buried surface area, pitch angle, shape complementarity, tilt angle, and Z distance showed little to no correlation with the CDR:pMHC PAE Score.

These results indicate that AF3’s predictive ability of T cell antigen specificity is based on the geometric orientation of the TCR relative to the pMHC, rather than any biochemical interactions the model can make. This suggests the model is not understanding the key drivers of antigen specificity [31], and may be a limitation for the model to fully understand the rules governing T cell antigen specificity.

### Improved standardisation of AlphaFold 3 derived binding scores

An important consideration for using AF3 to make predictions about T cell antigen specificity is the confounding factors that influence the confidence scores. By design, these scores aim to estimate the quality of the model’s structure predictions, and thus, each TCR and pMHC will have an intrinsic confidence, regardless if its binding a cognate partner. In their work, Bradley [13] introduces background interactions to normalise complex predictions and reduce the effects of these confounders. Further, the nature of the PAE scores creates skewed probability density distributions at protein-protein interfaces where higher confidences (lower PAE scores) are more present (see Figure 5A and B). Here we introduce the PAE Aggregator Score, a learned score for predicting TCR:pMHC binding from the PAE information to reduce the confounding factors.

**Figure 5:**
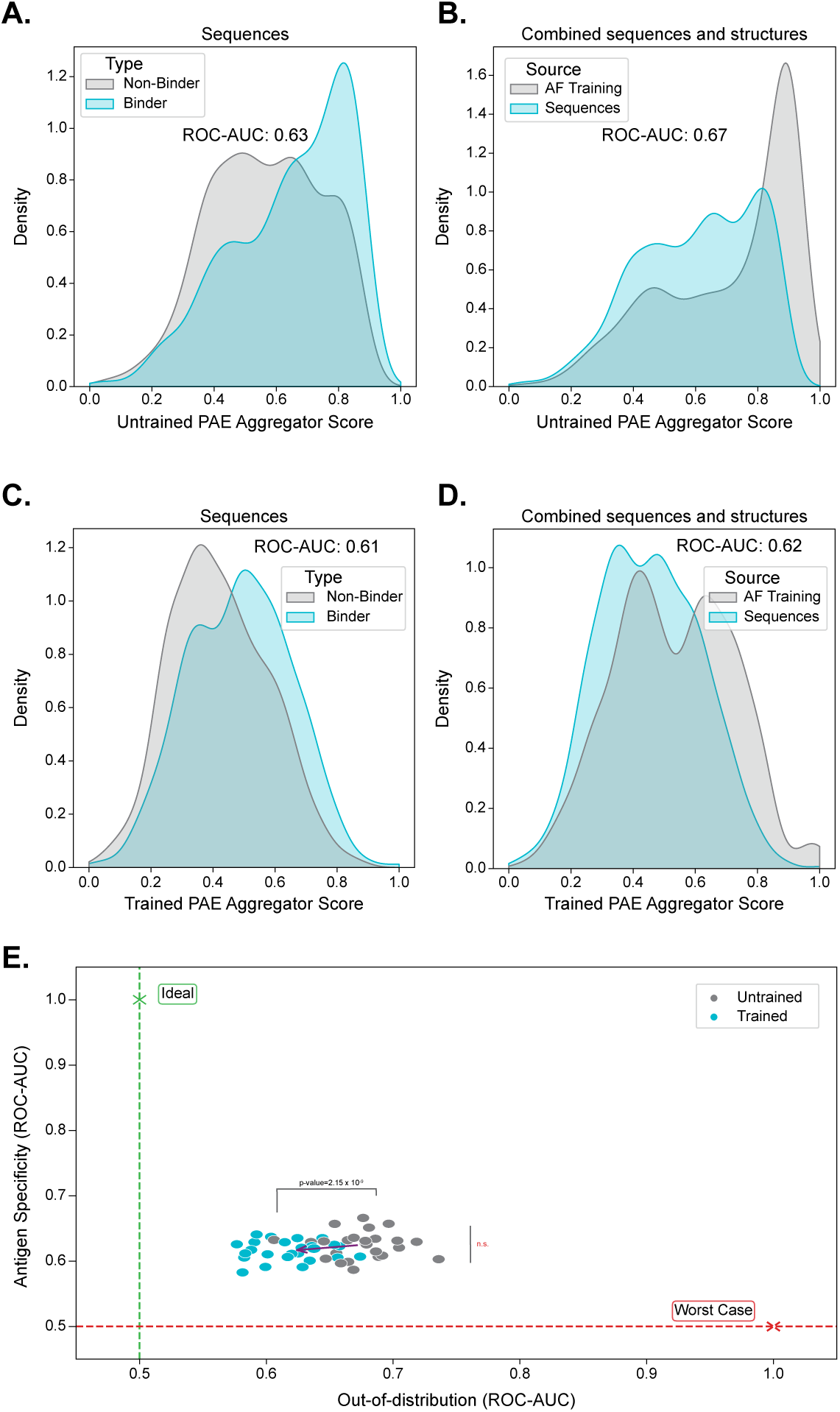
Improvement of specificity scores from using the PAE Aggregator model. **A.** Probability density distribution of the untrained PAE Aggregator Scores (same as CDR:pMHC PAE score normalised) comparing the scores ability to differentiate TCR:pMHC specificity in the 1500 TCR:pMHC complex sequences. **B.** Probability density distribution of the untrained PAE Aggregator Scores (same as CDR:pMHC PAE score normalised), illustrating the difference between predicting novel sequences and sequences taken from structures within the AF3 training cut-off, regardless of binding status. **C.** Probability density distribution of the trained PAE Aggregator Scores comparing the scores ability to differentiate TCR:pMHC specificity in the 1500 TCR:pMHC complex sequences. **D.** Probability density distribution of the trained PAE Aggregator Scores for novel sequences and sequences taken from structures within the AF3 training cut-off. **E.** Bootstrap predictions of untrained and trained PAE aggregator scores for predicting antigen specificity (y-axis) or novel sequences versus training structure-derived sequences (x-axis).

As described in “PAE Aggregator Binding Score”, the PAE Aggregator Score learns the relative importance of each CDR-peptide/MHC residue pair when averaging the CDR:pMHC PAE Scores to create a binding score for a given TCR:pMHC complex. The QQ-plots of Figure S6 show that after training, the PAE Aggregator Scores are much more normally distributed (R^2^ of 0.937 and 0.974 before training, compared to an R^2^ of 0.995 and 0.990 after training). This is visually apparent when comparing Figure 5 panels A and B to panels C and D.

Figure 5 also shows that the PAE Aggregator Score removes some of the confounding effects in the AF3-based binding scores. To simulate this, we compare the different binding scores’ ability to predict T cell antigen specificity versus the potential intrinsic effects of higher confidence TCR:pMHC complexes. For the antigen specificity prediction, we use our dataset of 1500 pMHC:pMHC sequences, and to explore the effect of higher confidence complexes, we supplement our 1500 sequences with 330 sequences (115 binders and 115 shuffled non-binders) that come from structures within the AF3 training cut-off. These additional sequences represent intrinsically higher confidence TCRs and pMHCs, as AF3 will have most likely seen them during training and thus score them more confidently. Before training, the PAE Aggregator Score, which is the same as the previously used CDR:pMHC PAE Score after normalisation, shows a large effect based on whether the complex sequence was from the sequence dataset or from the AF3 training structures. A ROC-AUC score of 0.67 separates these two sub-distributions. However, after training, the separations drop to a ROC-AUC of 0.62 (Figure 5 panel B versus panel D). The distributions also become bimodal, indicating that the learned binding score better captures the antigen specificity information instead of the intrinsic TCR or pMHC confidence. However, the performance of predicting antigen specificity stays roughly the same; a ROC-AUC of 0.63 is reached before training, and a ROC-AUC of 0.61 is reached after training (Figure 5 panel A versus panel C).

We statistically quantify all of these effects by conducting a bootstrap analysis. Shown in Figure 5E, there is statistically significant improvement after training in reducing the effect of intrinsically different TCR:pMHC confidences. A two-sided t-test with an adjusted significance level of 0.025 yielded a p-value of 2.15 *×* 10^−9^ that was deemed significant. In contrast, the ability of the scores to predict T cell antigen specificity showed no significance after the same statistical test was performed.

Our PAE Aggregator Score provides a TCR:pMHC complex-specific binding score based on the AF3 confidence scores. The learned score improves the normality of predictions and reduces the confounding effects of intrinsic confidence scores.

### AlphaFold 3 captures distinct binding modes of sequence repertoires

To further investigate AF3’s ability to decode antigen specificity, we used the model to understand the features of TCRs with common specificity for a single antigen. In their work, Dash et al. [32] produced a dataset of TCR repertoires from different patients binding to several epitopes to investigate TCR common specificity. Dash et al. [32] and Mayer and Callan [33] conducted clustering analyses on the 92 TCRs that recognise the BMLF epitope in this dataset that was derived from Epstein-Barr Virus. The clustering analyses revealed that there were several dominant clusters (encompassing multiple donors), with internally similar sequences (< 5 Levenshtein distance) but differing sequences between clusters [33] (see Figure 6A). There were also single TCRs that were distinct from any major cluster. It was hypothesised that the differing sequence clusters represented different binding modes of TCRs to the same pMHC. We sought to determine whether the structures predicted by AF3 validated this hypothesis.

**Figure 6:**
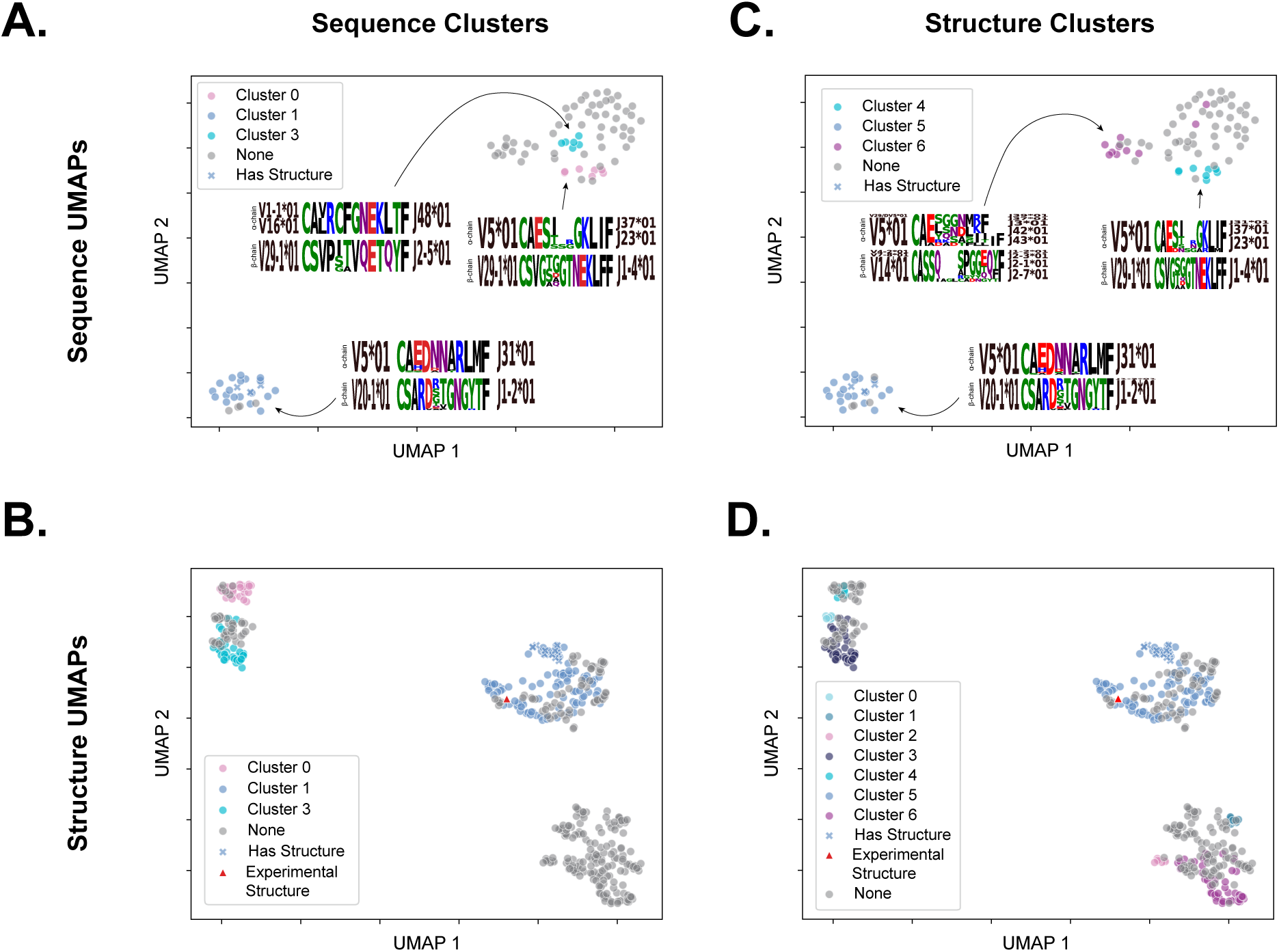
AlphaFold captures sequence-level clusters in distinct binding modes. **A.** uniform manifold approximation and projection (UMAP) of the TCR sequences reacting to the BMLF epitope from the Dash et al. [32] study. The dominant sequence clusters are coloured, and their motifs are annotated. For three sequences, a structure is available (PDB ID 3o4l); these have been annotated as crosses. **B.** UMAP of the TCR AF3 structures reacting to the BMLF epitope with the clusters derived from the CDR3 sequences coloured. The experimental structure (PDB ID 3o4l) is also included and shown as a red triangle. **C.** UMAP of the TCR sequences reacting to the BMLF epitope with the clusters derived from AF3 structures coloured. **D.** UMAP of the TCR AF3 structures reacting to the BMLF epitope with clusters derived from AF3 structures coloured.

We performed the same clustering as Mayer and Callan [33], taking the combined Levenshtein distance of the CDR3α and CDR3β and clustering TCRs with a combined distance of 5 or less and that grouped at least 3 TCRs. The TCR Levenshtein distance space is visualised in Figure 6A, with the sequence clusters coloured and annotated with their motif. It is apparent that the clustered sequences group closely together in the UMAP visualisation of the sequence space. We then predicted the structure of the 92 TCRs in complex with the BMLF epitope and HLA-A*02:01 from the Dash et al. [32] dataset with AF3 and created numerical representations using the same structural features described in “Parameterising TCR:pMHC complexes”. Figure 6B shows the structural space in a UMAP representation, overlaying the same sequence-derived clusters. Visually, it is apparent that the sequence clusters correspond to distinct regions of predicted structure space. This provides evidence that the different sequence clusters do correspond to different binding modes. For three of the sequences, a structure has been determined that is within the AF3 training cut-off (PDB ID 3o4l). We included both the predicted structures of these sequences as well as the resolved crystal structure as validation of the structural quality. We see that the predicted and experimental structures group closely together in the UMAP space, supporting the quality of the AF3 predictions.

Next, we aimed to cluster the BMLF-specific TCRs by their structural features, and visualise the correspondence in sequence space. We used hierarchical density-based spatial clustering of applications with noise (HDBSCAN) to create clusters derived from the structural properties of the TCR:pMHC complexes. We determined the optimal minimum cluster size as eight based on the elbow plot, shown in Figure S7. The structural clusters are visualised in relation to the sequence space and structural space in Figure 6C and D, respectively. For visualisation in sequence space, we did the same filtering of clusters with fewer than 3 TCRs. The structural clusters have some differences from the sequence-based clustering in the TCRs they group in sequence space (Adjusted Rand Index of 0.54 and V-measure of 0.64). While two clusters remain similar (sequence cluster 0-structure cluster 4 and sequence cluster 1-structure cluster 5), sequence cluster 3 and structure cluster 6 are quite dissimilar. The structure cluster has a more diverse CDR3 motif and is dominated by TRAV5*01 and TRBV14*01, whereas the sequence cluster is made up of TRAV1–1*01, TRAV16*01, and TRBV29–1*01. The structure cluster is also made up of a variety of TRAJ and TRBJ genes, while the sequence cluster is restricted to TRAJ48*01 and TRBJ2–5*01. The structure clusters encompass slightly more TCRs compared to the sequence clusters, clustering 43% and 38% of the 92 TCR:pMHC complexes, respectively.

Overall, the structural featurisation and clustering approaches taken with AF3 show comparable results to the sequence-based clustering. This provides additional support that the differences in sequence-based clustering are driven by alternative binding modes employed by the TCRs to bind the BMLF epitope.

### AlphaFold 3 has the sensitivity to detect specificity-disrupting point mutations

We aimed to determine the sensitivity of AF3 at predicting antigen specificity by assessing its ability to detect disruptive single amino acid mutations. In their recent work, Jones et al. [34] describe the effects of mutations on the 572G TCR’s ability to bind coeliac antigens. The TCR cross-reacts to two closely related coeliac antigens presented by HLA-DQ2.5 (HLA-DQA1*05:01 and HLA-DQB1*02:01): the α1 epitope (PFPQPELPY) and the ω1 (PFPQPEQPF) epitope. Structures are available within the AF3 training cutoff for the antigens presented by HLA-DQ2.5, but there are no structures of the 572G TCR within the cutoff. Through surface plasmon resonance (SPR) experiments, Jones et al. [34] found that the TCR has higher affinity for the α1 epitope (K_D_ of 6.1 µM) over the ω1 epitope (K_D_ of 37.8 µM). They identified the central tryptophan (W) and aspartate (D) of the CDR3β as being key drivers of the TCR’s specificity through crystallography, and confirmed using SPR that mutating either of these residues disrupts the binding, while mutating other residues in the loop preserves some affinity. We tested AF3’s ability to elucidate these results using our PAE Aggregator Score. We use a normalised affinity value, dividing the minimum affinity value (6.1 µM) by the affinity values so that the highest affinity is represented as 1 and the maximum K_D_ value is 0.

Shown in Figure 7A, after predicting the wild-type 572G2 TCR in complex with both the α1 and ω1 epitopes, the difference in binding score reflects the reported affinity difference between these two antigens. Performing a one-sided Wilcoxon rank-sum test shows that the α1 complex binding scores are significantly higher than the ω1 complex at a significance level of 0.05, achieving a p-value of 3.97 *×* 10^−3^. However, although both complexes can be classified as “Medium” predictions using the CAPRI evaluation of DockQ scores, a two-sided Wilcoxon rank-sum test shows that the α1 ternary complex is of higher quality than the ω1 complex (considering a significance level of 0.05 and a p-value of 7.94 *×* 10^−3^; see Figure 7B). We also see that both the normalised affinity and DockQ scores are correlated with the Binding Scores, but the DockQ correlation is stronger than the affinity (R^2^ = 0.85 versus R^2^ = 0.11; see Figure 7C and D). This further illustrates the confounding factors in AF3’s binding score discussed in “Improved standardisation of AlphaFold 3 derived binding scores” and makes the results ambiguous.

**Figure 7:**
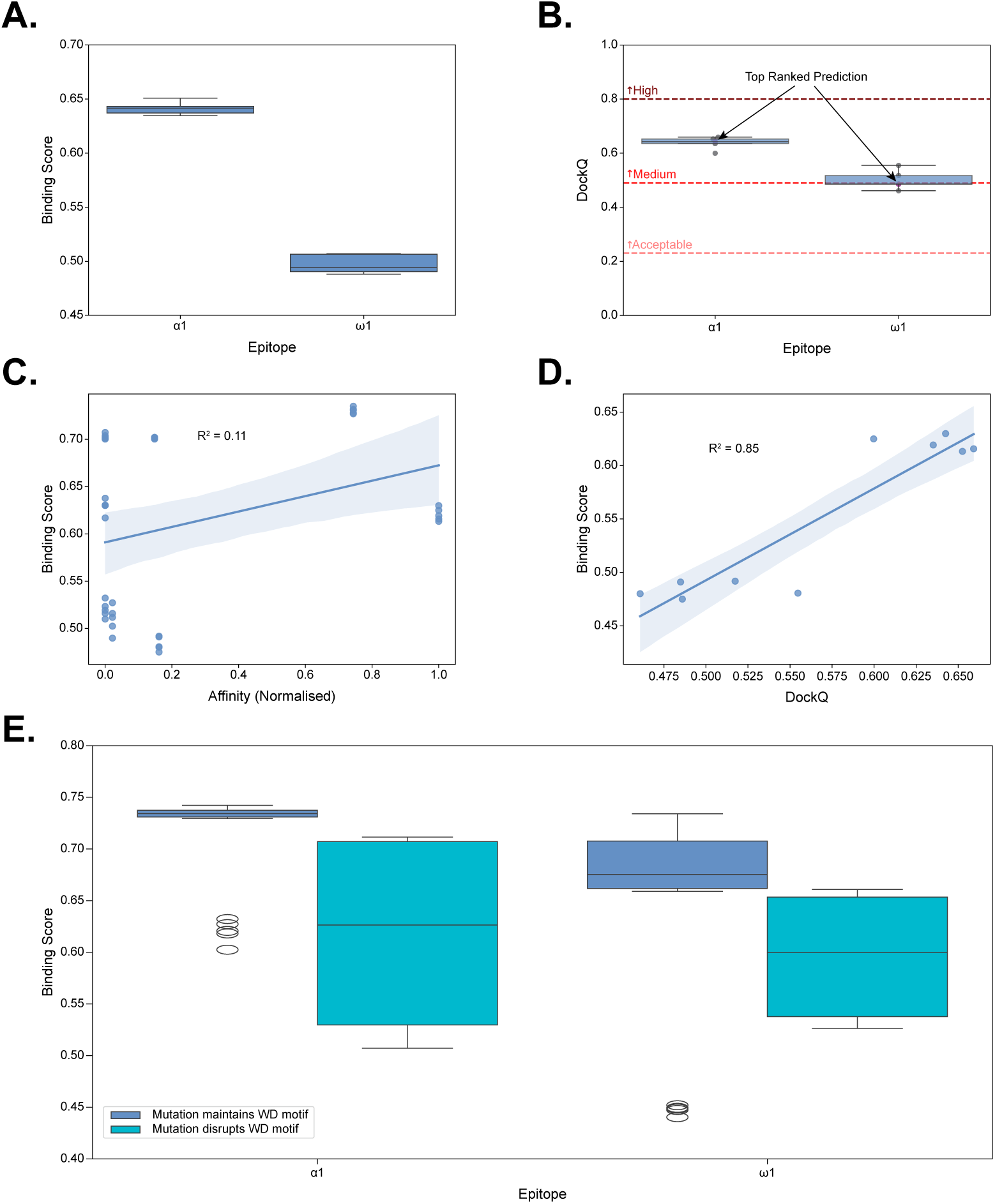
AlphaFold-derived binding scores capture TCR mutations that are known to disrupt antigen specificity. **A.** Comparison of the AF3-derived binding scores for the 572G2 TCR binding to the α1 and ω1 epitopes. **B.** Comparison of the DockQ scores of the 572G2 TCR binding to the α1 and ω1 epitopes compared with the ground truth structures. The CAPRI grading of DockQ scores is annotated. **C.** Correlation between the AF3-derived binding scores and the normalised affinity of the 572G2 TCR and alanine-scanning mutants binding to α1 and ω1 epitopes. **D.** Correlation of the DockQ scores to binding scores for the 572G2 TCR binding to the α1 and ω1 epitopes. **E.** Comparison of the binding scores for the alanine-scanning mutants of the 572G2 TCR binding to α1 and ω1 epitopes. The mutants are stratified by epitope and also whether they disrupt the tryptophan (W)-aspartate (D) motif previously reported to be key for binding both of these epitopes.

Turning to the mutational scanning experiments, AF3 is successful at detecting the disrupting mutations. In Figure 7E, we show the difference in binding scores between mutations that affect the key tryptophan or aspartate residues and those that affect other parts of the CDR3. A two-factor ANOVA test shows that there is a significant difference at a significance level of 0.05 in the binding score of the mutations that disrupt the WD motif and those that do not (p-value = 2.03 *×* 10^−4^), as well as confirming that there is a significant difference in between the binding scores of the α1 and ω1 epitopes (p-value = 1.62 *×* 10^−3^). There were no significant interaction effects reported. The changes in binding score correspond to ROC-AUCs of 0.92 for the α1 epitope and 0.83 for the ω1 epitope. These results are in line with task-specific models, such as P-TEAM, that are trained on these types of mutational scanning experiments directly and require data on a particular TCR-epitope pair [35].

The success of AF3 at detecting these disrupting mutations reinforces its ability to predict T cell antigen specificity. However, its sensitivity may not be enough to conduct full *in silico* optimisation campaigns of a TCR to a target.

## Discussion

In this work, we explore AF3’s ability to predict T cell antigen specificity in novel settings. After creating and validating a high-throughput pipeline for TCR:pMHC complex predictions, we benchmarked AF3 against the open-licence methods used in IMMREP25 [15] and showed that it was the top-performing model at both predicting T cell antigen specificity and TCR:pMHC complex structures. We evaluated the impact of the input signals to the model, establishing the importance of the MSAs, in particular the unpaired MSAs, for TCR:pMHC predictions. We then looked at the underlying correlates of its performance and found that the confidence scores correlated strongly with the placement of a TCR:pMHC and not any biochemical interactions. We further developed the PAE Aggregator Score as a TCR:pMHC specific binding score that reduces the confounding factors in the AlphaFold confidence scores. Finally, we investigated AF3’s ability to cluster antigens and detect disrupting mutations, and found that it has the specificity and sensitivity to do both well.

While AF3 has shown great promise at solving the unseen epitope challenge, the problem is still a long way from being solved. The ROC-AUC performance of AF3 reached a maximum of *≈* 0.6. Although this is better than previous models, which have near-random performance (ROC-AUC *≈* 0.5), it does not achieve good enough discrimination of binders and non-binders to remove the need for experimental validation or to be considered for use in clinical settings. Performance would need to be upwards of ROC-AUC *≈* 0.9 to match the ability of current experimental assays. Further, although we and others have improved the throughput of AF3 for predicting TCR:pMHC complexes, it is still computationally expensive to perform large screens of TCR:pMHC complexes. It took around 5 days to predict the structures of our sampled 1500 sequences on a single NVIDIA RTX A6000 with 48 GB of memory; to run the entire dataset without sub-sampling on this hardware, it would have taken over 250 days. To fully solve the unseen epitope challenge, we suspect that data creation of more TCR:pMHC structures for model training, more TCR:pMHC sequences for MSA generation, and advances in model design for increased throughput will be essential.

Another limitation of using structure prediction-based strategies for decoding T cell antigen specificity is identifying the most representative prediction from the predicted ensemble. Fromm et al. [36] demonstrate that in the related task of predicting antibody-antigen complexes, increasing the number of samples improves results, but the ranking score helps very little in being able to identify the top prediction. We see in this work that selecting only the top-ranked prediction from the ranking score did little to improve the performance on any task. We also see this in our case study of the 572G TCR binding the α1 and ω1 coeliac antigens: the top-ranked predictions based on the AlphaFold ranking score do not correspond to the highest DockQ scores. As a result, we did not explore increasing the number of seeds or samples to attempt to sample more conformational space and get better results in this work.

This study, as well as most others in the field, suffers from a lack of true negative binding. In this work, we employ a shuffled-epitope approach (see “Collecting sequence data”) to generate suspected non-binding TCR:pMHC pairs; however, this relies on the assumption that non-reported interactions in the data will most likely be negative. The alternatives would be to either use reported non-binders, such as those in the IEDB [37], or sample TCRs from healthy cohorts as non-reactive TCRs and pair those to epitopes in our dataset as negatives. However, other works have shown that both of these alternatives introduce larger sources of bias than the shuffled epitope approach [38]. Therefore, the benchmarks of predicting T cell antigen specificity presented in this work must be taken with this context.

Of note in the case study of the 572G TCR binding both the α1 and ω1 epitopes, a mutation in the V gene encoded region at the N-terminus of the CDR from a serine to alanine (CASS… to CASA…) was found as an outlier (see Figure 7E). The mutation conferred a similar binding score penalty as the disruptive tryptophan and aspartate mutations. In Jones et al. [34], this mutation was used as a negative control and showed that mutants with it maintained specificity, which is in opposition to what is presented here. However, *≈* 70% of the CDR3βs of available TCR structures begin with CASS [29]. We therefore suspect that AF3 has (incorrectly) learned this as essential for TCR structures, and modifying this region results in a loss of confidence for any structure prediction. These types of considerations are important to bear in mind while interpreting predictions based on AF3.

Further, it must be taken into account that our analysis of AF3’s ability to detect point mutations was conducted on two specific cases: the 572G TCR mutants binding to the α1 and ω1 epitopes and the c259 TCR binding the NY-ESO-1 peptide mutants. These results outlined the ability and limitations of AF3 in these specific cases, and while the results were in agreement, they are not necessarily representative of the wider ability of AF3. A broader study of AF3 correlation to affinity measure could be conducted using a dataset such as ATLAS [39], but full TCR:pMHC sequence information is not always present, and consideration would need to be made across different affinity measurement techniques.

A major advancement introduced by AF3 is predicting all-atom representations of biomolecular structures to encompass small molecules, lipids, and RNA/DNA. This ability could enable the prediction of antigen specificity for unconventional TCRs [40]. These TCRs bind non-canonical peptides and non-protein antigens, which most previous sequence-based methods would have little support for (except TITAN [8]).

An area in which we see the potential for further improvement of these models is building more informative MSAs. Although our results showed the value MSAs give to TCR:pMHC predictions, we believe this is the area with the biggest opportunity for improvement. Currently, the MSAs created by AlphaFold, in particular the paired MSA representation, rely on evolutionary information to identify co-occurring residue mutations. In contrast, the stochasticity of the TCR:pMHC interface, which is essential for immune recognition, is not represented in the MSAs. If the conserved amino acid substitutions of TCR:pMHC binding could be captured in the input for the model, similar to the ideas presented in Postovskaya et al. [41] and Pyo et al. [42], it could enable better modelling of these interactions. However, it would be important to balance these signals with the information that AlphaFold has accrued about general protein-protein interactions, which seems to power its success at decoding TCR specificity.

## Methods

### Data curation

#### Collecting sequence data

We curated a dataset of 13,962 unique pairs of TCRs binding to pMHCs with complete sequence information by collating several publicly available datasets. We downloaded sequences from the VDJdb [43] on January 10th, 2025 using the selection criteria of “CDR3 – Species: Human, Monkey, Mouse”, “CDR3 – Gene (chain): TRA, TRB”, “MHC – Class: MHCI, MHCII”, “Meta – Assay Type: Multimer sorting, Culture-based, Other”, “Meta – Sequencing: Sanger, High-throughput, Single-cell”, and “Meta – Spurious CDR3: Include non-canonical, Include unmapped V/J”. Next, we downloaded all the sequences from McPas-TCR [44] on January 10th, 2025. We then downloaded sequences from the immune epitope database (IEDB) [37] on January 14th, 2025 using the selection criteria of“Assay: T cell, MHC Ligand, Outcome – Positive”, “Epitope: Any”, “MHC Restriction: Any”, “Host: Any”, “Disease: Any”, and “Reference: Any”. Finally, we downloaded the ITRAP dataset [45] on January 14th, 2025.

From these sequence datasets, we kept only the entries that had full paired α- and β-chain information (V gene, CDR3, and J gene), a peptide antigen, and an MHC-class I allele listed. We also removed entries in the VDJdb that came from a 10x Genomics assay^1^, as this 10x study has been reported to contain a high number of false positives [45]. We replaced these entries with the filtered version provided by the ITRAP dataset.

We then ensured that all gene codes and CDR3 regions were standardised using the tidytcells Python package [46]. Following that, we used the Stitchr Python package [47] to create full-length TCR sequences from the V gene code, CDR3, J gene code, and species information. Next, we extracted the CDR1, CDR2, and CDR3 amino acid sequences using ANARCI [48]. To create full-length MHC sequences from the allele codes, we downloaded the data from the IMGT/HLA and IMGT/MHC databases and matched the allele codes of the TCR:pMHC complexes to the canonical sequences in the IMGT data [49]. We then isolated MHC pseudo-sequence representations from the full-length MHC sequences (see “Defining TCR-MHC pseudo representation positions” for details). Finally, we deduplicated these results based on the six CDRs, peptide, and MHC pseudo-sequence to create the final dataset of 13,962 unique binding TCR:pMHCs.

To generate negative data of non-interacting TCR:pMHC pairs, we employed a shuffled epitope approach. For every peptide in our positive binding dataset, we randomly sampled five times the number of positive binding TCRs from all other TCRs in the dataset that were reported to bind different peptides. In cases where there were fewer than five times the number of alternative TCRs available, we included all of them. After this process, we ended up with a dataset of 83,772 TCR:pMHCs where approximately one-fifth (13,962) were reported binders from the literature and the remainder (69,810) were potential negatives generated through our shuffling procedure.

We partitioned our data into five splits of roughly equal size to create cross-validation folds for downstream modelling tasks. We grouped the TCR:pMHC pairs by the peptide and the MHC involved in the interaction. These groups containing differing numbers of cognate TCRs were distributed as equally as possible into five folds. This ensured that there was no data leakage from the pMHC pairing information.

After collating the sequence data, we downsampled the sequences into samples of 50 and 1500 TCR:pMHC pairs. We ensured that an equal number of data points were drawn from each cross-validation fold, and an equal number of positive and negative pairings were selected.

#### Collecting structure data

We curated a dataset of 251 TCR:pMHC complexes from the STCRDab [29] which we used throughout this analysis. We downloaded all entries in the STCRDab on September 5th, 2024. From these, we selected all αβTCRs binding to pMHC-Is or pMHC-IIs presenting peptide antigens. We kept all structures with a resolution better than 3.50 Å. We discarded structures that were missing residues in key domains of the CDRs loops or peptide. In cases where there were missing residues in the TCR framework region or MHC antigen binding domain, we used MODELLER [50] to remodel the missing residues, using the original PDB file as the only template and only allowing the missing residues’ position to be optimised by the algorithm. We then isolated TCR:pMHC complexes from the asymmetric units of the PDB files and extracted the sequences of their CDRs and peptides. We added MHC allele information by querying the PDB IDs of our isolated structures on histo.fyi [51]. We removed complexes with redundant CDR, peptide, and MHC allele information that also had less than 2 Å overall root mean square deviation (RMSD) to the selected TCR:pMHC complex. Finally, we removed all hetero atoms from the structure files.

In their work, Shi et al. [24] identified TCR:pMHC structures in the PDB with little sequence identity to the structures within the AF3 training cutoff. For our hold-out set to evaluate structure prediction quality, we used our selected structures that were also in this set of sequence-dissimilar complexes. This created a set of 21 TCR:pMHC structures for evaluation with the following PDB ids: 6zkw, 7dzm, 7na5, 7ndq, 7ndt, 7nme, 7ow5, 7pbe, 7phr, 7q9b, 7rk7, 7t2b, 7t2d, 7z50, 8d5q, 8dnt, 8gom, 8gvi, 8i5c, 8i5d, and 8shi.

### Defining TCR-MHC pseudo representation positions

In this work, we use TCR-MHC pseudo representations, a representation of MHC molecules that is mainly concerned with TCR binding [52, 53]. These representations take inspiration from the MHC pseudo-sequences used in the peptide presentation predictor, NetMHCpan [54]. To calculate the TCR-pMHC pseudo-representation, we count all of the contacts occurring between heavy atoms on the TCR CDRs and the MHC at a distance of less than 5 Å in our structure dataset. We calculate the frequency of these contacts for each international immunogenetics information system (IMGT) position on the MHC molecules, normalising the MHC class-Is and MHCs class-IIs separately before combining based on IMGT position and renormalising. Finally, we filter out any positions that make up less than 1% of contacts. From this, we identify the following MHC IMGT positions as the TCR-MHC pseudo representation:

- α1-helix: 62, 63, 65, 66, 68, 69, 70, 72, 73, 75, 76, 79
- α2-helix: 58, 61A, 62, 63, 65, 66, 67, 69, 70, 72, 72A, 73, 76, 77

This representation can be used to create MHC pseudo-sequences that are specific to TCR binding, or can be used to represent other information at these positions on MHC structures.

### Accelerated data pipeline for TCR:pMHC structure prediction

We developed a pipeline to maximise TCR:pMHC complex prediction throughput using AlphaFold. The “data” and “folding” stages of AlphaFold inference were split apart. All sequences for the MHC chains, TCR α-chains, and TCR β-chains were deduplicated individually and precomputed. The data pipeline was not run for the peptide sequences as these do not return MSA or templates. The databases provided by TCRmodel2 [23] were used in place of the default AF3 databases. These were generated by Yin et al. [23] by running a handful of representative TCRs and pMHCs through the AF2 databases and deduplicating the results. They are a small subset of the original databases, which allows for faster scanning of sequences in the MSA construction phase. From these MSAs and templates, the folding pipeline for a TCR:pMHC complex can be run by combining the relevant MSAs and adding an empty MSA for the peptide.

### Running AlphaFold 3 (and other models) folding pipeline

The folding pipeline was run with default settings for all models except Boltz-2, where the number of diffusion samples was increased to 5 to match the other models. A random seed of ‘1’ was used for all folding pipelines, and the default 200 diffusion steps were used for AF3, Chai-1, and Boltz-2. AF3 predictions were run with the default ten recycles everywhere, except when comparing to other models; here, three recycles were used to match the other models. AF2 was run using the structure relaxing setting turned on, as this has been reported to improve its performance [21].

### Deriving a binding score from the structure prediction confidence scores

#### CDR:pMHC PAE Score

We derived a binding score based on the AF3 PAE confidence measures in a similar fashion to Bradley [13] that we call the CDR:pMHC PAE Score. For each TCR:pMHC complex, we take the mean PAE value for all residues in the TCR CDRs, relative to all residues in the peptide and TCR-MHC pseudo representation. This is presented as follows:

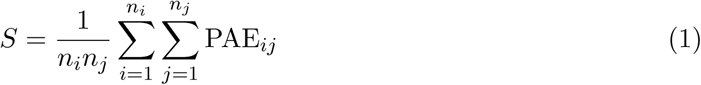

Where *n_i_* is the number of CDR residues and *n_j_*is the number of residues in the peptide and TCR-MHC pseudo representation.

#### PAE Aggregator Binding Score

Expanding on the CDR:pMHC PAE Score, we created a binding score we termed the PAE Aggregator Binding Score by learning the relative weights of the previous CDR:pMHC PAE score. In the previous formulation, the CDR:pMHC PAE score relied on an equally weighted average of the relevant PAE positions. Here, we train a simple machine learning model to learn the relative importance of each PAE residue pair.

As a training dataset, we use the predicted structures of our 1500 TCR:pMHC complex sequences, as well as the predicted structures of sequences derived from structures in our structure dataset that are before the AF3 training cut-off (2021–09–30) [16]. For the 115 structures from our structural dataset before the AlphaFold training cut-off, we employed a shuffled negative approach on the sequences (as described in “Collecting sequence data”) to create 115 negatives. Additionally, the structures were annotated with cross-validation folds following the peptide-MHC driven folds from the sequence data. In cases where the structures did not share a peptide or MHC with any of the sequences, they were randomly allocated to 1 of the 5 cross-validation folds. This results in a dataset of 9150 predicted structures: 1500 from the sampled sequences, 330 sequences from structures, and 5 predicted structures per sequence. For all predicted structures, we extract the PAE entries corresponding to our CDR:pMHC PAE Score, and added blank rows and columns as padding to achieve a length of 8 for the CDRs 1 and 2s, a length of 24 for the CDR 3s, and a length of 12 for the peptides. The padding is done to centre the entities. For even-length entities, the blank rows or columns are added to either side of the entity; for odd-length entities, the extra insertion is done on the preceding entity. Finally, the PAE values are normalised with the following definition:

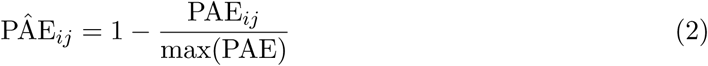

so that the highest confidence values are 1 and the lowest confidence values are zero. The padding is also represented as zero.

To learn the relative importance of PAE residues, the CDR:pMHC PAE Score was reformulated as follows:

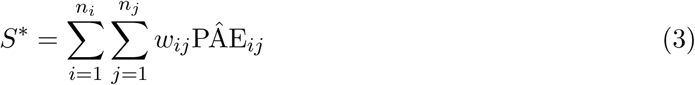

Where *w_ij_* represents a learnable parameter with an initial value of 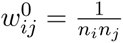. An additional bound was placed on the learned score to keep it between 0 and 1:

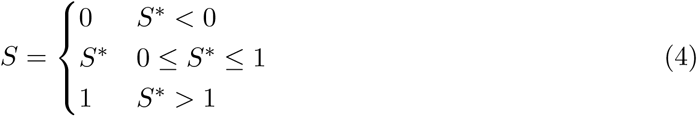

With this formulation, the untrained model is the same as the original CDR:pMHC PAE Score normalised. To learn the weights, the model was trained to predict the binding status (‘0’ for non-binders or ‘1’ for binders) for the 9150 predicted structures using a five-fold cross-validation strategy. For each holdout fold, the model was trained in 15 epochs on the remaining data with a mini-batch size of 32 and a learning rate of 1 *×* 10^−5^ using a binary cross-entropy loss function.

### Bootstrapping for antigen specificity prediction assessment

To evaluate the structure prediction models’ ability to predict antigen specificity, we used a bootstrapped ROC-AUC assessment. For our dataset of 50 TCR:pMHC complexes that was used to evaluate the MSA generation strategy, we conducted 15 trials, and in each trial we randomly sampled 20 complexes and evaluated the ROC-AUC performance of predicting our labels of binding versus non-binding complexes. For our dataset of 1500 TCR:pMHC complexes, which was used in the ablation studies and benchmarking of similar models, we conducted 25 bootstrap trials and sampled 500 TCR:pMHC complexes in each trial. When evaluating our trained PAE Aggregator Binding Score, we used the hold-out data and the version of the model trained on the remaining TCR:pMHC complexes for each cross-validation fold to create predictions for each entry in our dataset. We then performed the same bootstrapping with 25 trials and 500 sampled points in each trial to assess the performance of the learned model.

### Measuring the DockQ score as a metric of structure quality

We used DockQ [55] to evaluate the structure prediction models’ ability to predict TCR:pMHC complex structures. DockQ combines the fraction of native contacts recovered, the interface RMSD, and the ligand (pMHC) RMSD. This was done using the implementation in the STCRpy package [30].

### Correlating TCR:pMHC complex features to AF3 confidence scores

#### Parameterising TCR:pMHC complexes

To parametrise both experimental and predicted TCR:pMHC complex structures, we created a set of 12 features. These included the pitch angle, tilt angle, roll angle, scanning angle, buried surface area, shape complementarity, X distance, Y distance, Z distance, number of interactions, number of interaction types, and contact maps. To calculate the pitch angle and scanning angle, we used the docking geometry calculator from STCRpy in ‘Cys’ mode [30]. This method aligns the TCR:pMHC complex to a reference MHC molecule, assigning a coordinate system using vectors defined by the antigen binding domain helices and the reference MHC centre-of-mass. The TCR is then given a vector and centre-of-mass based on the conserved cysteine residues in the TCR framework region. We similarly used the values of the TCR centre-of-mass over the reference MHC centre-of-mass as the X, Y, and Z distance features. To calculate the tilt and roll angle, we used the same coordinate system defined by the reference MHC molecule and TCR vector. The tilt angle is taken as the angle the TCR vector makes off of the Z-axis in the YZ-plane. This differs from the pitch angle, which is calculated as the angle directly off the Z-axis. Similarly, the roll angle is taken as the angle the TCR vector makes to the Z-axis in the XZ-plane.

Both the number of interactions and the number of interaction types are taken by using the protein-ligand interaction profiler (PLIP) [56] through STCRpy [30] between the TCR and peptide. The number of interactions is a count of all interactions in the TCR:pMHC complex, whereas the number of interaction types is the count of distinct interactions made, including hydrogen bonds, hydrophobic interactions, salt bridges, or π-stacks.

The buried surface area of TCR:pMHC complexes was computed using the Shrake-Rupley molecular surface area algorithm implemented in Biopython [57]. The buried area was calculated as the difference between the combined individual TCR and pMHC surface areas, and the TCR:pMHC surface area. The default 1.40 Å probe radius and 100-point resolution were used.

Shape complementarity was computed using the following Python package: https://github.com/t-whalley/SCASA. The package scores two surfaces’ shape complementarity between 0 and 1 by randomly sampling points on each surface and measuring the dot product of the adjacent surface normals.

Finally, contact maps were calculated by measuring the probability of equivalent residues with heavy atoms within 5 Å of each other between the TCR and pMHC. Equivalent residues were determined by centre numbering the CDRs and peptides. For odd-length entities, the residue in the centre was denoted as position 0, with everything towards the N-terminus being negatively numbered and everything towards the C-terminus positively numbered. For even-length entities, position 0 was denoted as the left-centre residue so that more positive numbers extended to the C-terminus. For the MHC molecules, equivalent residues were determined using IMGT numbering of the MHC.

#### Selecting structural features and fitting multiple linear regression model

To develop statistical associations of our 12 structural features with the AF3 confidence scores, we processed the features and fit a multiple linear regression model. The probability density distributions of the scanning angles, tilt angles, roll angles, X distances, Y distances, Z distances, number of interactions, shape complementarities, and buried surface areas were all right-skewed. To correct this, we applied a quantile transform using the scikit-learn package’s implementation with 100 quantiles, targeting normal distributions for these features [58]. We also conducted a correlation analysis between our structural features and removed the scanning angles and roll angles, as these had high correlation (*>* 0.5) to the contact maps and tilt angle, respectively. With this processed selected set of features, we fit a multiple linear regression model with an intercept between them and the CDR:pMHC PAE score using the statsmodels Python package [59].

#### Bootstrap anaylsis of structural correlates to AF3 confidence scores

A bootstrap analysis was conducted to ascertain the robustness of structural feature correlations to the AF3 confidence scores. First, outliers were removed using a Mahalanobis distance of 0.95; points with a Mahalanobis distance greater than this from the distribution of selected structural features and CDR:pMHC PAE scores were removed. Next, 100 trials were performed, randomly sampling 1000 points each trial and fitting a multiple linear regression in the same way as described above in “Selecting structural features and fitting multiple linear regression model”. In each trial, the fitted parameters of the selected structural features were subjected to a one-sided t-test, assessing whether the parameter was negative (indicating that higher confidence structures- lower PAE scores- fit with canonical TCR binding). In cases where the p-value was less than a significance level of 0.05, the result was noted as significant.

## Acknowledgments

This work was supported by funding from the UK Medical Research Council grant number MC UU 12010/3 to H.K., the UK Medical Research Council grant number MC UU 00008 to B.Mc., and an ARISE Fellowship from the European Union’s Horizon 2020 Research and Innovation Programme under the Marie Sk-lodowska-Curie grant agreement number 945405 to C.T. We acknowledge the support of the Natural Sciences and Engineering Research Council of Canada (NSERC).

We would also like to thank the developers of PyMOL, Pandas, pingouin, NumPy, Matplotlib, Scikit-Learn, SciPy, and seaborn for their contributions to the open-source community. These tools and packages were used to conduct the analysis and generate the figures for this work.

## Author contributions

**B.Mc.**: Investigation, Formal analysis, Software, Writing – Original Draft. **A.E.M.**: Investigation, Writing – Review & Editing. **I.I.**: Methodology, Writing – Review & Editing. **C.J.T.**: Supervision. **N.L.L.G**: Methodology, Resources. **J.R.**: Methodology, Resources. **C.M.D.**: Supervision, Writing – Review & Editing. **H.K.**: Supervision, Writing – Review & Editing.

## Data and code availability

The data generated and used in this manuscript are available at https://doi.org/10.5281/zenodo.21242412. All of the code and notebooks used to conduct this analysis can be found at https://github.com/benjiemc/tcr-pmhc-predicted-structures.

## Footnotes

1 https://www.10xgenomics.com/resources/application-notes/a-new-way-of-exploring-immunity-linking-highly-multiplexed-antigen-recognition-to-immune-repertoire-and-phenotype/#

## References

[1] Stephen J. Turner et al. “Structural Determinants of T-cell Receptor Bias in Immunity”. In: Nature Reviews Immunology 6.12 (Dec. 1, 2006), pp. 883–894. issn: 1474-1733, 1474-1741. doi: 10.1038/nri1977. url: https://www.nature.com/articles/nri1977 (visited on 04/03/2023).

[2] Veronika Zarnitsyna et al. “Estimating the Diversity, Completeness, and Cross-Reactivity of the T Cell Repertoire”. In: Frontiers in Immunology 4 (Dec. 26, 2013). issn: 1664-3224. doi: 10.3389/fimmu.2013.00485. url: https://www.frontiersin.org/journals/immunology/articles/10.3389/fimmu.2013.00485/full (visited on 04/01/2026).

[3] Steven G. E. Marsh et al. “Nomenclature for Factors of the HLA System, 2026”. In: HLA 107.3 (2026), e70595. issn: 2059-2310. doi: 10.1111/tan.70595. url: https://onlinelibrary.wiley.com/doi/abs/10.1111/tan.70595 (visited on 06/04/2026).

[4] Chloe H. Lee et al. “Predicting Cross-Reactivity and Antigen Specificity of T Cell Receptors”. In: Frontiers in Immunology 11 (Oct. 22, 2020). issn: 1664-3224. doi: 10.3389/fimmu.2020.565096. url: https://www.frontiersin.org/journals/immunology/articles/10.3389/fimmu.2020.565096/full (visited on 04/01/2026).

[5] Dan Hudson et al. “Can We Predict T Cell Specificity with Digital Biology and Machine Learning?” In: Nature Reviews Immunology 23.8 (Aug. 2023), pp. 511–521. issn: 1474-1741. doi: 10.1038/s41577-023-00835-3. url: https://www.nature.com/articles/s41577-023-00835-3 (visited on 11/06/2024).

[6] Alessandro Montemurro et al. “NetTCR-2.0 Enables Accurate Prediction of TCR-peptide Binding by Using Paired TCRα and β Sequence Data”. In: Communications Biology 4.1 (1 Sept. 10, 2021), pp. 1–13. issn: 2399-3642. doi: 10.1038/s42003-021-02610-3. url: https://www.nature.com/articles/s42003-021-02610-3 (visited on 10/21/2022).

[7] Pieter Moris et al. “Current Challenges for Unseen-Epitope TCR Interaction Prediction and a New Perspective Derived from Image Classification”. In: Briefings in Bioinformatics 22.4 (July 1, 2021), bbaa318. issn: 1477-4054. doi: 10.1093/bib/bbaa318. url: 10.1093/bib/bbaa318 (visited on 10/31/2022).

[8] Anna Weber, Jannis Born, and María Rodriguez Martínez. “TITAN: T-cell Receptor Specificity Prediction with Bimodal Attention Networks”. In: Bioinformatics 37 (Supplement 1 July 1, 2021), pp. i237–i244. issn: 1367-4803. doi: 10.1093/bioinformatics/btab294. url: 10.1093/bioinformatics/btab294 (visited on 10/24/2022).

[9] Zahra S. Ghoreyshi and Jason T. George. “Quantitative Approaches for Decoding the Specificity of the Human T Cell Repertoire”. In: Frontiers in Immunology 14 (Sept. 7, 2023). issn: 1664-3224. doi: 10.3389/fimmu.2023.1228873. url: https://www.frontiersin.org/journals/immunology/articles/10.3389/fimmu.2023.1228873/full (visited on 04/01/2026).

[10] Filippo Grazioli et al. “On TCR Binding Predictors Failing to Generalize to Unseen Peptides”. In: Frontiers in Immunology 13 (Oct. 21, 2022), p. 1014256. issn: 1664-3224. doi: 10.3389/fimmu.2022.1014256. PMID: 36341448. url: https://www.ncbi.nlm.nih.gov/pmc/articles/PMC9634250/ (visited on 11/10/2022).

[11] Lihua Deng et al. “Performance Comparison of TCR-pMHC Prediction Tools Reveals a Strong Data Dependency”. In: Frontiers in Immunology 14 (Apr. 18, 2023). issn: 1664-3224. doi: 10.3389/fimmu.2023.1128326. url: https://www.frontiersin.org/journals/immunology/articles/10.3389/fimmu.2023.1128326/full (visited on 04/02/2024).

[12] Morten Nielsen et al. “Lessons Learned from the IMMREP23 TCR-epitope Prediction Challenge”. In: ImmunoInformatics 16 (Dec. 1, 2024), p. 100045. issn: 2667-1190. doi: 10.1016/j.immuno.2024.100045. url: https://www.sciencedirect.com/science/article/pii/S2667119024000156 (visited on 10/07/2024).

[13] Philip Bradley. “Structure-Based Prediction of T Cell Receptor:Peptide-MHC Interactions”. In: eLife 12 (Jan. 20, 2023). Ed. by Michael L Dustin, Tadatsugu Taniguchi, and Michael L Dustin, e82813. issn: 2050-084X. doi: 10.7554/eLife.82813. url: 10.7554/eLife.82813 (visited on 02/21/2023).

[14] Benjamin McMaster et al. “Can AlphaFold’s Breakthrough in Protein Structure Help Decode the Fundamental Principles of Adaptive Cellular Immunity?” In: Nature Methods 21.5 (May 2024), pp. 766–776. issn: 1548-7105. doi: 10.1038/s41592-024-02240-7. url: https://www.nature.com/articles/s41592-024-02240-7 (visited on 11/06/2024).

[15] Eve Richardson et al. IMMREP25: Unseen Peptides. Apr. 1, 2026. doi: 10.64898/2026.03.30.715276. url: https://www.biorxiv.org/content/10.64898/2026.03.30.715276v1 (visited on 04/09/2026). Pre-published.

[16] Josh Abramson et al. “Accurate Structure Prediction of Biomolecular Interactions with AlphaFold 3”. In: Nature 630.8016 (June 2024), pp. 493–500. issn: 1476-4687. doi: 10.1038/s41586-024-07487-w. url: https://www.nature.com/articles/s41586-024-07487-w (visited on 08/11/2025).

[17] Chai Discovery et al. Chai-1: Decoding the Molecular Interactions of Life. Oct. 15, 2024. doi: 10.1101/2024.10.10.615955. url: https://www.biorxiv.org/content/10.1101/2024.10.10.615955v2 (visited on 10/08/2025). Pre-published.

[18] Jeremy Wohlwend et al. Boltz-1 Democratizing Biomolecular Interaction Modeling. May 6, 2025. doi: 10.1101/2024.11.19.624167. url: https://www.biorxiv.org/content/10.1101/2024.11.19.624167v4 (visited on 10/08/2025). Pre-published.

[19] Saro Passaro et al. Boltz-2: Towards Accurate and Efficient Binding Affinity Prediction. June 18, 2025. doi: 10.1101/2025.06.14.659707. url: https://www.biorxiv.org/content/10.1101/2025.06.14.659707v1 (visited on 04/01/2026). Pre-published.

[20] John Jumper et al. “Highly Accurate Protein Structure Prediction with AlphaFold”. In: Nature 596.7873 (7873 Aug. 2021), pp. 583–589. issn: 1476-4687. doi: 10.1038/s41586-021-03819-2. url: https://www.nature.com/articles/s41586-021-03819-2 (visited on 10/20/2022).

[21] Brennan Abanades et al. “ImmuneBuilder: Deep-Learning Models for Predicting the Structures of Immune Proteins”. In: Communications Biology 6.1 (1 May 29, 2023), pp. 1–8. issn: 2399-3642. doi: 10.1038/s42003-023-04927-7. url: https://www.nature.com/articles/s42003-023-04927-7 (visited on 05/30/2023).

[22] Nele P. Quast et al. “T-Cell Receptor Structures and Predictive Models Reveal Comparable Alpha and Beta Chain Structural Diversity despite Differing Genetic Complexity”. In: Communications Biology 8.1 (Mar. 4, 2025), p. 362. issn: 2399-3642. doi: 10.1038/s42003-025-07708-6. url: https://www.nature.com/articles/s42003-025-07708-6 (visited on 02/05/2026).

[23] Rui Yin et al. “TCRmodel2: High-Resolution Modeling of T Cell Receptor Recognition Using Deep Learning”. In: Nucleic Acids Research (May 4, 2023), gkad356. issn: 0305-1048. doi: 10.1093/nar/gkad356. url: 10.1093/n ar/gkad356 (visited on 05/10/2023).

[24] Yudan Shi, Jerry M. Parks, and Jeremy C. Smith. “Comparative Analysis of TCR and TCR-pMHC Complex Structure Prediction Tools”. In: Journal of Chemical Information and Modeling 65.13 (July 14, 2025), pp. 7156–7173. issn: 1549-9596. doi: 10.1021/acs.jcim.5c00298. url: 10.1021/acs.jcim.5c00298 (visited on 09/05/2025).

[25] Jiadong Lu et al. “Benchmarking TCR–pMHC Structure Prediction: A Unified Evaluation and CDR3-based Functional Insights”. In: Briefings in Bioinformatics 27.3 (May 1, 2026), bbag289. issn: 1477-4054. doi: 10.1093/bib/bbag289. url: 10.1093/bib/bbag289 (visited on 07/07/2026).

[26] Martin Steinegger and Johannes Söding. “MMseqs2 Enables Sensitive Protein Sequence Searching for the Analysis of Massive Data Sets”. In: Nature Biotechnology 35.11 (Nov. 2017), pp. 1026–1028. issn: 1546-1696. doi: 10.1038/nbt.3988. url: https://www.nature.com/articles/nbt.3988 (visited on 01/07/2026).

[27] Milot Mirdita et al. “ColabFold: Making Protein Folding Accessible to All”. In: Nature Methods 19.6 (2022), pp. 679–682. issn: 1548-7091. doi: 10.1038/s41592-022-01488-1. PMID: 35637307. url: https://pmc.ncbi.nlm.nih.gov/articles/PMC9184281/ (visited on 01/07/2026).

[28] Sebastian N. Deleuran and Morten Nielsen. “NetTCR-struc, a Structure Driven Approach for Prediction of TCR-pMHC Interactions”. In: Frontiers in Immunology 16 (July 17, 2025). issn: 1664-3224. doi: 10.3389/fimmu.2025.1616328. url: https://www.frontiersin.org/journals/immunology/articles/10.3389/fimmu.2025.1616328/full (visited on 08/14/2025).

[29] Jinwoo Leem et al. “STCRDab: The Structural T-cell Receptor Database”. In: Nucleic Acids Research 46.D1 (Jan. 4, 2018), pp. D406–D412. issn: 0305-1048. doi: 10.1093/nar/gkx971. url: 10.1093/nar/gkx971 (visited on 12/07/2022).

[30] Nele P Quast, Charlotte M Deane, and Matthew I J Raybould. “STCRpy: A Software Suite for T-cell Receptor Structure Parsing, Interaction Profiling, and Machine Learning Dataset Preparation”. In: Bioinformatics 41.10 (Oct. 1, 2025), btaf566. issn: 1367-4811. doi: 10.1093/bioinformatics/btaf566. url: 10.1093/bioinformatics/btaf566 (visited on 11/20/2025).

[31] Lars Kjer-Nielsen et al. “A Structural Basis for the Selection of Dominant Alpha-beta T Cell Receptors in Antiviral Immunity”. In: Immunity 18.1 (Jan. 2003), pp. 53–64. issn: 1074-7613. doi: 10.1016/s1074-7613(02)00513-7. PMID: 12530975.

[32] Pradyot Dash et al. “Quantifiable Predictive Features Define Epitope Specific T Cell Receptor Repertoires”. In: Nature 547.7661 (July 6, 2017), pp. 89–93. issn: 0028-0836. doi: 10.1038/nature22383. PMID: 28636592. url: https://www.ncbi.nlm.nih.gov/pmc/articles/PMC5616171/ (visited on 10/21/2022).

[33] Andreas Mayer and Curtis G. Callan. “Measures of Epitope Binding Degeneracy from T Cell Receptor Repertoires”. In: Proceedings of the National Academy of Sciences 120.4 (Jan. 24, 2023), e2213264120. doi: 10.1073/pnas.2213264120. url: https://www.pnas.org/doi/abs/10.1073/pnas.2213264120 (visited on 04/03/2025).

[34] Claerwen M. Jones et al. “Gliadin-Specific CD4+ T Cells Exhibit Functional Diversity Independent of TCR Usage or Epitope Specificity”. In: Submitted Unknown.Unknown (Jan. 1, 2026), Unknown. issn: Unknown. doi: Unknown. url: Unknown.

[35] Felix Drost et al. “Predicting T Cell Receptor Functionality against Mutant Epitopes”. In: Cell Genomics 4.9 (Sept. 11, 2024), p. 100634. issn: 2666-979X. doi: 10.1016/j.xgen.2024.100634. url: https://www.sciencedirect.com/science/article/pii/S2666979X24002386 (visited on 06/22/2026).

[36] Samuel Fromm, Marko Ludaic, and Arne Elofsson. “Evaluating Deep Learning Based Structure Prediction Methods on Antibody–Antigen Complexes”. In: Bioinformatics 42.4 (Apr. 1, 2026), btag136. issn: 1367-4811. doi: 10.1093/bioinformatics/btag136. url: 10.1093/bioinformatics/btag136 (visited on 04/20/2026).

[37] Randi Vita et al. “The Immune Epitope Database (IEDB): 2024 Update”. In: Nucleic Acids Research 53.D1 (Jan. 6, 2025), pp. D436–D443. issn: 1362-4962. doi: 10.1093/nar/gkae1092. url: 10.1093/nar/gkae1092 (visited on 01/08/2026).

[38] Ceder Dens et al. “The Pitfalls of Negative Data Bias for the T-cell Epitope Specificity Challenge”. In: Nature Machine Intelligence 5.10 (Oct. 2023), pp. 1060–1062. issn: 2522-5839. doi: 10.1038/s42256-023-00727-0. url: https://www.nature.com/articles/s42256-023-00727-0 (visited on 04/23/2025).

[39] Tyler Borrman et al. “ATLAS: A Database Linking Binding Affinities with Structures for Wild-Type and Mutant TCR-pMHC Complexes”. In: Proteins: Structure, Function, and Bioinformatics 85.5 (2017), pp. 908–916. issn: 1097-0134. doi: 10.1002/prot.25260. url: https://onlinelibrary.wiley.com/doi/abs/10.1002/prot.25260 (visited on 03/13/2023).

[40] Dale I. Godfrey et al. “The Burgeoning Family of Unconventional T Cells”. In: Nature Immunology 16.11 (11 Nov. 2015), pp. 1114–1123. issn: 1529-2916. doi: 10.1038/ni.3298. url: https://www.nature.com/articles/ni.3298 (visited on 12/14/2022).

[41] Anna Postovskaya et al. “tcrBLOSUM: An Amino Acid Substitution Matrix for Sensitive Alignment of Distant Epitope-Specific TCRs”. In: Briefings in Bioinformatics 26.1 (Nov. 22, 2024), bbae602. issn: 1467-5463. doi: 10.1093/bib/bbae602. PMID: 39576224. url: https://pmc.ncbi.nlm.nih.gov/articles/PMC11583439/ (visited on 04/20/2026).

[42] Andrew G. T. Pyo et al. “Data-Driven Discovery of Biophysical T Cell Receptor Cospecificity Rules”. In: PRX Life 3.3 (July 15, 2025), p. 033005. issn: 2835-8279. doi: 10.1103/14j1-wrh5. url: https://link.aps.org/doi/10.1103/14j1-wrh5 (visited on 10/08/2025).

[43] Mikhail Shugay et al. “VDJdb: A Curated Database of T-cell Receptor Sequences with Known Antigen Specificity”. In: Nucleic Acids Research 46.D1 (Jan. 4, 2018), pp. D419–D427. issn: 0305-1048. doi: 10.1093/nar/gkx760. url: 10.1093/nar/gkx760 (visited on 10/21/2022).

[44] Nili Tickotsky et al. “McPAS-TCR: A Manually Curated Catalogue of Pathology-Associated T Cell Receptor Sequences”. In: Bioinformatics 33.18 (Sept. 15, 2017), pp. 2924–2929. issn: 1367-4803. doi: 10.1093/bioinformatics/btx286. url: 10.1093/bioinformatics/btx286 (visited on 10/21/2022).

[45] Helle Rus Povlsen et al. “Improved T Cell Receptor Antigen Pairing through Data-Driven Filtering of Sequencing Information from Single Cells”. In: eLife 12 (May 3, 2023). Ed. by K Christopher Garcia, Tadatsugu Taniguchi, and Michael E Birnbaum, e81810. issn: 2050-084X. doi: 10.7554/eLife.81810. url: 10.7554/eLife.81810 (visited on 01/08/2026).

[46] Yuta Nagano and Benjamin Chain. “Tidytcells: Standardizer for TR/MH Nomenclature”. In: Frontiers in Immunology 14 (Oct. 25, 2023). issn: 1664-3224. doi: 10.3389/fimmu.2023.1276106. url: https://www.frontiersin.org/journals/immunology/articles/10.3389/fimmu.2023.1276106/full (visited on 01/08/2026).

[47] James M Heather et al. “Stitchr: Stitching Coding TCR Nucleotide Sequences from V/J/CDR3 Information”. In: Nucleic Acids Research 50.12 (July 8, 2022), e68. issn: 0305-1048. doi: 10.1093/nar/gkac190. url: 10.1093/nar/gkac190 (visited on 01/08/2026).

[48] James Dunbar and Charlotte M. Deane. “ANARCI: Antigen Receptor Numbering and Receptor Classification”. In: Bioinformatics 32.2 (Jan. 15, 2016), pp. 298–300. issn: 1367-4803. doi: 10.1093/bioinformatics/btv552. url: 10.1093/bioinformatics/btv552 (visited on 11/09/2023).

[49] James Robinson et al. “IMGT/HLA and IMGT/MHC: Sequence Databases for the Study of the Major Histocompatibility Complex”. In: Nucleic Acids Research 31.1 (Jan. 1, 2003), pp. 311–314. issn: 1362-4962. doi: 10.1093/nar/gkg070. PMID: 12520010.

[50] Andrej Šali and Tom L. Blundell. “Comparative Protein Modelling by Satisfaction of Spatial Restraints”. In: Journal of Molecular Biology 234.3 (Dec. 5, 1993), pp. 779–815. issn: 0022-2836. doi: 10.1006/jmbi.1993.1626. url: https://www.sciencedirect.com/science/article/pii/S0022283683716268 (visited on 01/14/2026).

[51] Christopher J. Thorpe. Histo.Fyi — An Interactive Exploration of the Structure and Function of MHC Molecules. url: https://www.histo.fyi/ (visited on 09/05/2024).

[52] Benjamin McMaster et al. “Quantifying Conformational Changes in the TCR:pMHC-I Binding Interface”. In: Frontiers in Immunology 15 (Dec. 2, 2024). issn: 1664-3224. doi: 10.3389/fimmu.2024.1491656. url: https://www.frontiersin.org/journals/immunology/articles/10.3389/fimmu.2024.1491656/full (visited on 12/02/2024).

[53] Sagar Gupta and Nikolaos G. Sgourakis. “A Structure-Guided Approach to Predict MHC-I Restriction of T Cell Receptors for Public Antigens”. In: Structure 33.9 (Sept. 4, 2025), 1614–1623.e2. issn: 0969-2126. doi: 10.1016/j.str.2025.06.011. url: https://www.sciencedirect.com/science/article/pii/S096921262500245X (visited on 03/03/2026).

[54] Ilka Hoof et al. “NetMHCpan, a Method for MHC Class I Binding Prediction beyond Humans”. In: Immunogenetics 61.1 (Jan. 2009), pp. 1–13. issn: 0093-7711. doi: 10.1007/s00251-008-0341-z. PMID: 19002680. url: https://pmc.ncbi.nlm.nih.gov/articles/PMC3319061/ (visited on 02/12/2026).

[55] Claudio Mirabello and Björn Wallner. “DockQ v2: Improved Automatic Quality Measure for Protein Multimers, Nucleic Acids, and Small Molecules”. In: (). url: 10.1093/bioinformatics/btae586 (visited on 04/15/2026).

[56] Melissa F Adasme, et al. “PLIP 2021: Expanding the Scope of the Protein–Ligand Interaction Profiler to DNA and RNA”. In: Nucleic Acids Research 49.W1 (July 2, 2021), W530–W534. issn: 0305-1048. doi: 10.1093/nar/gkab294. url: 10.1093/nar/gkab294 (visited on 04/09/2026).

[57] Peter J. A. Cock et al. “Biopython: Freely Available Python Tools for Computational Molecular Biology and Bioinformatics”. In: Bioinformatics 25.11 (June 1, 2009), pp. 1422–1423. issn: 1367-4803. doi: 10.1093/bioinformatics/btp163. url: 10.1093/bioinformatics/btp163 (visited on 04/09/2026).

[58] Fabian Pedregosa et al. “Scikit-Learn: Machine Learning in Python”. In: Journal of Machine Learning Research 12.85 (2011), pp. 2825–2830. issn: 1533-7928. url: http://jmlr.org/papers/v12/pedregosa11a.html (visited on 04/10/2026).

[59] Skipper Seabold and Josef Perktold. “Statsmodels: Econometric and Statistical Modeling with Python”. In: SciPy 2010 (May 1, 2010). doi: 10.25080/Majora-92bf1922-011. url: https://proceedings.scipy.org/articles/Majora-92bf1922-011 (visited on 04/10/2026).

