## Supplementary document for "Characterising AlphaFold 3’s ability to predict T cell antigen specificity"

### Supplementary Information for "Characterising AlphaFold 3's ability to predict T cell antigen specificity"

#### Further correlation of the PAE Aggregator Score to affinity

As the analysis of the 572G TCR binding to coeliac antigens focused on mutations made to the TCR, we conducted a complementary analysis of the c259 TCR binding to a mutational scanning library of peptide variants reported by Cabezas-Caballero et al. [2]. The c259 TCR is an engineered therapeutic TCR targeting the NY-ESO-1 peptide (SLLMWITQV). In their work, Cabezas-Caballero et al. perform a full mutational scan, swapping all peptide positions for the 19 alternative amino acids and conduct SPR experiments to measure their affinity. We took this library, predicted the TCR:pMHC structures using AlphaFold, and then compared our PAE Aggregator Score to the normalised affinities. The normalisation was done by scaling the affinity values so that the true binding affinity (0.701 µM) was 1, and everything over 100 µM was 0. In some cases, the affinity values were not reported because either the TCR did not bind the mutant peptide, or the peptide was not presented by the MHC (HLA-A*02:01). We used NetMHCpan version 4.2 [3] to identify any of the missing values that were not classed as a strong or weak binder and excluded them from the analysis. For those that were missing affinity values and were NetMHCpan binders, we gave them a normalised affinity value of 0.

Fitting a linear model between the non-zero affinities and our binding score yielded an $R^2$ of 0.08, indicating a weak but existing association between our binding score and

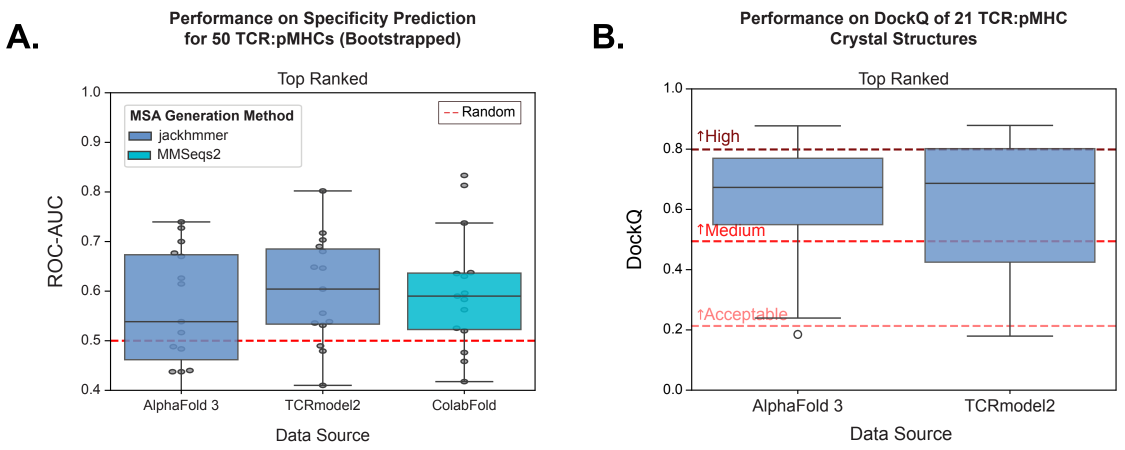


Figure S1: Effect of MSA strategy choice on performance considering only the top-ranked prediction for each complex. **A.** Boostrapped ROC-AUC performance predicting the specificity of 50 TCR:pMHC interactions. Random performance is annotated by the red dashed line. **B.** DockQ performance of predicting 21 AF3-holdout TCR:pMHC structures. The CAPRI grading of DockQ scores is annotated.

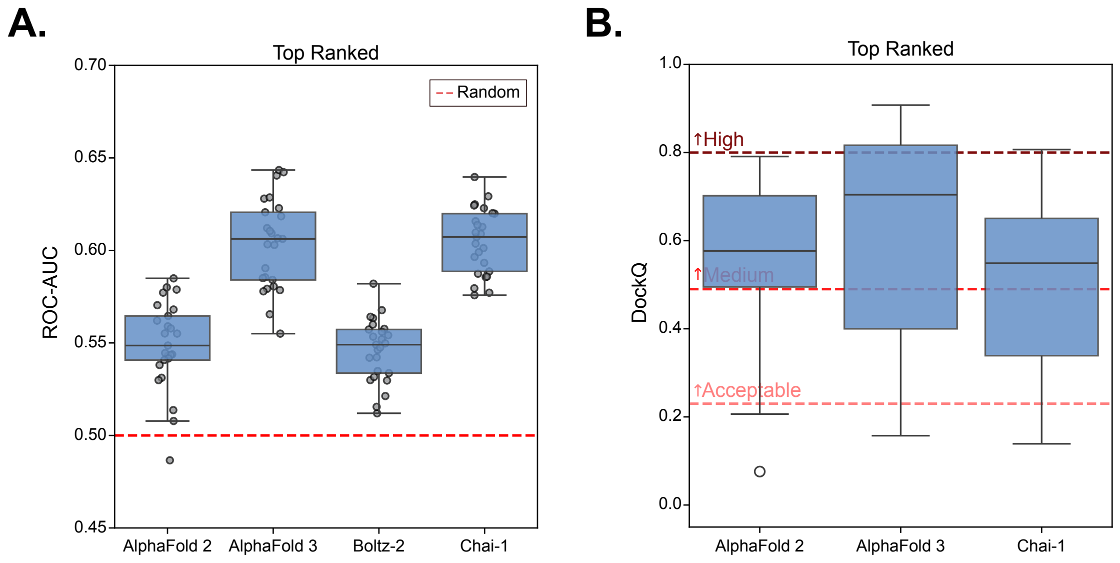


Figure S2: Benchmark of structure prediction models considering only the top-ranked prediction for each complex. **A.** ROC-AUC performance at predicting antigen specificity from 1500 TCR:pMHC. **B.** DockQ performance at predicting TCR:pMHC complex structures from the benchmark set. Boltz-2 was not included as only three structures were within the model's training cut-off The CAPRI grading of DockQ scores is annotated.

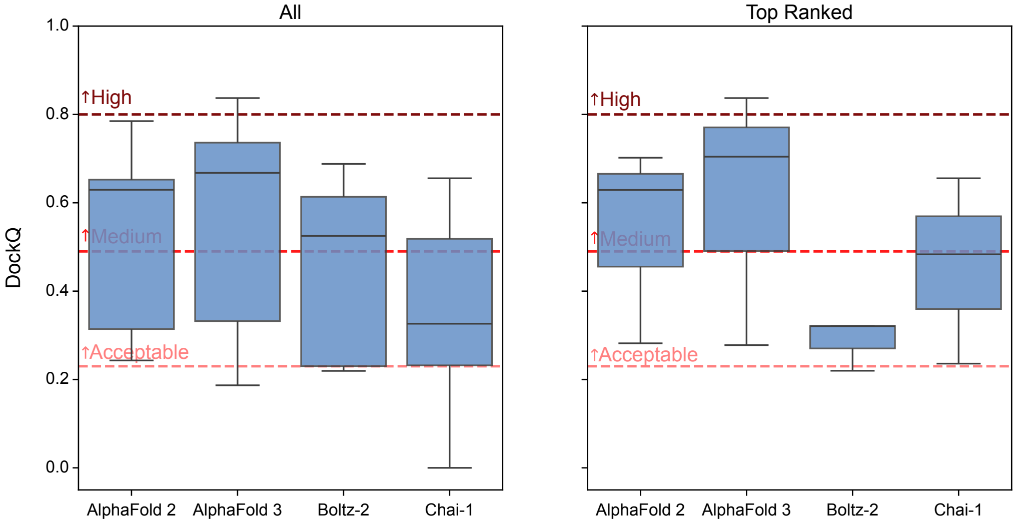


Figure S3: Benchmark of structure prediction models at predicting 8shi, 8i5c, and 8i5d (the three structures from the hold-out set outside of the Boltz-2 training cut-off date), considering (**A.**) all predictions, or (**B.**) only the top-ranked model prediction. The CAPRI grading of DockQ scores is annotated.

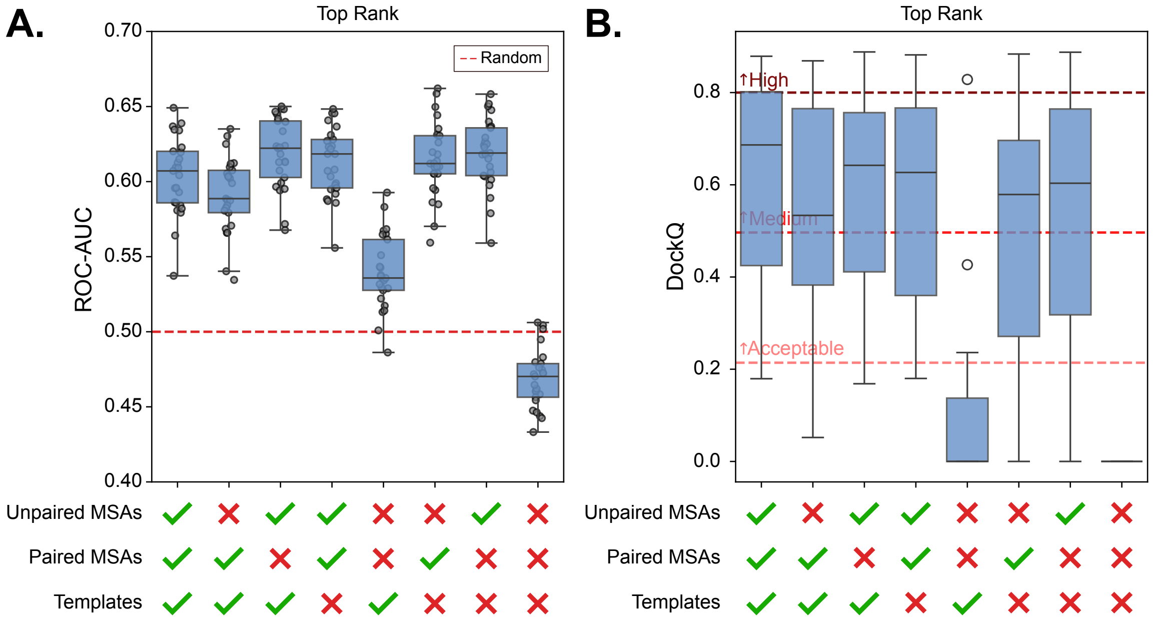


Figure S4: Effects of AF3 input channel ablations on performance considering only the top-ranked prediction for each complex. **A.** ROC-AUC performance at predicting T cell antigen specificity using 1500 TCR:pMHC interactions. **B.** DockQ performance at predicting TCR:pMHC complex structures. The CAPRI grading of DockQ scores is annotated.

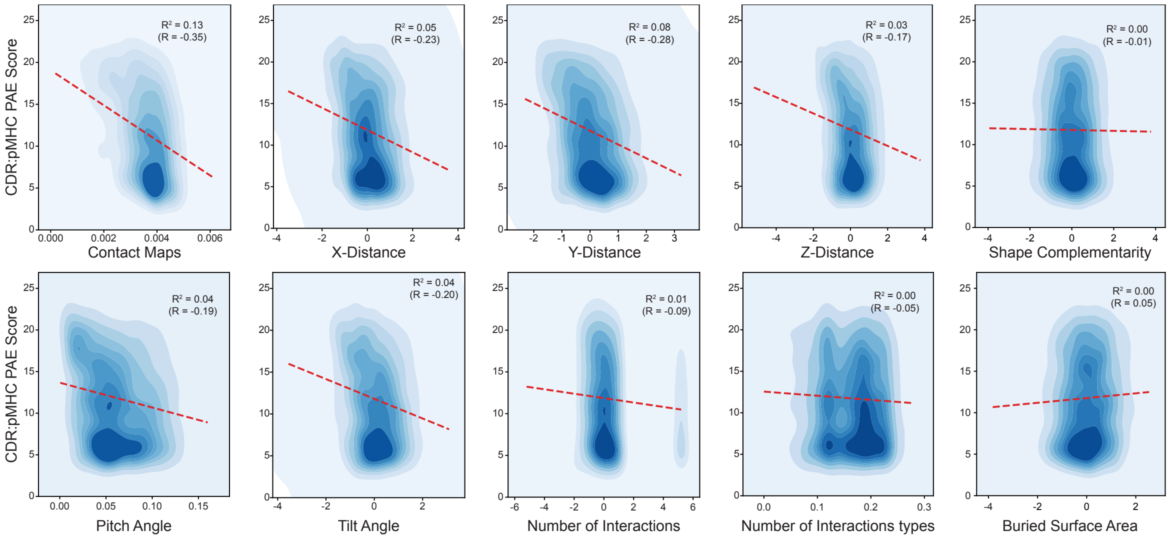


Figure S5: Individual linear regressions between each of the selected TCR:pMHC complex features and the CDR:pMHC PAE Binding Score.

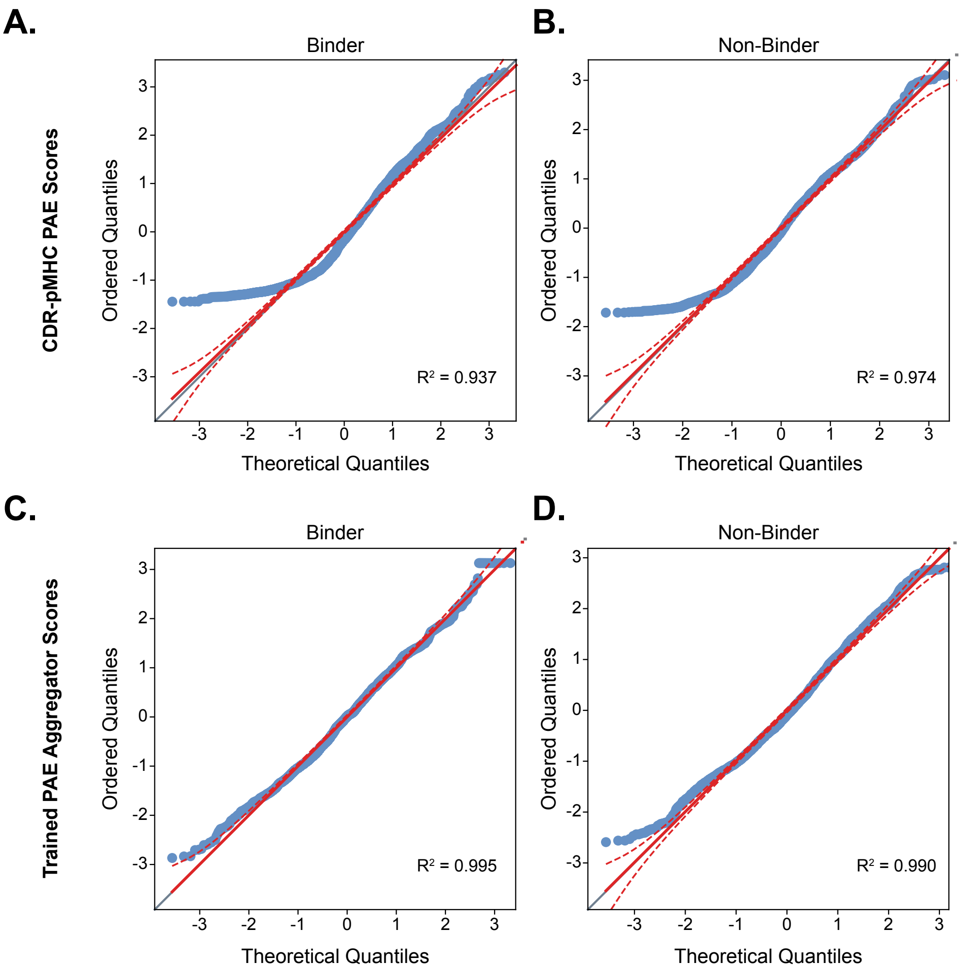


Figure S6: QQ-Plots comparing untrained and trained PAE Aggregator scores normality. **A.** Binding and **B.** non-Binding scores from original AF3 CDR:pMHC PAE score. **C.** Binding and **D.** non-Binding scores from trained AF3 PAE Aggregator Score.

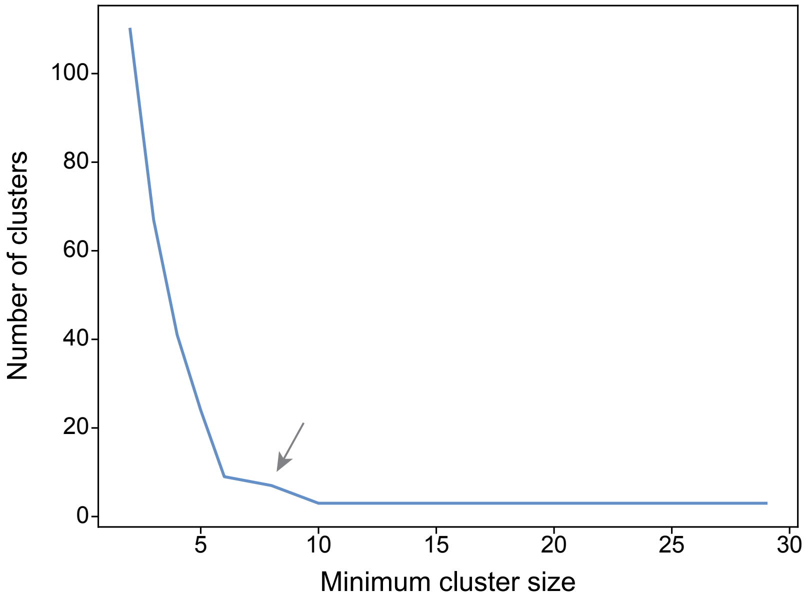


Figure S7: Elbow plot to determine the best minimum cluster size for the structural clusters of the TCRs specific to the BMLF epitope in the Dash et al. dataset. The elbow point was determined to be a minimum cluster size of 8, indicated by the arrow.

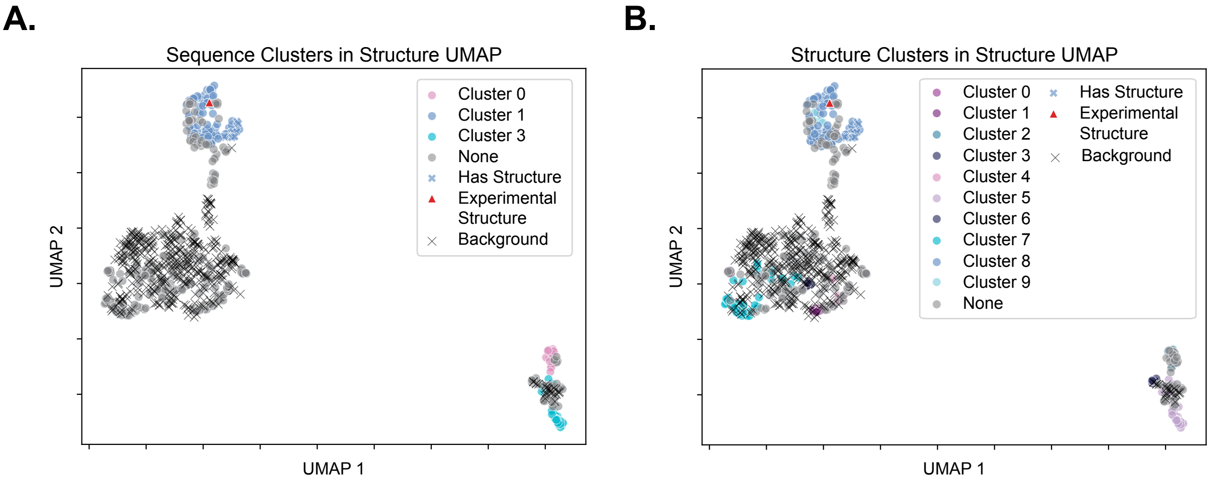


Figure S8: Clustering of TCRs binding the BMLF epitope in the Dash et al. dataset including background TCRs randomly sampled from the sequence dataset.

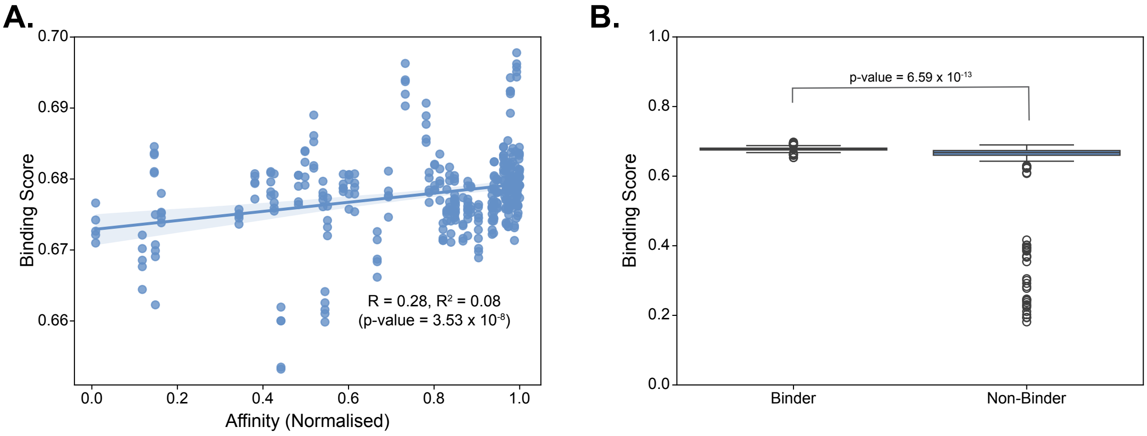


Figure S9: Comparison of PAE Aggregator Score with affinities of c259 TCR peptide mutational scan library from Cabezas-Caballero et al. [2] **A.** Regression of PAE Aggregator Scores with c259 TCR peptide mutant affinities **B.** Comparison of binding (< 100 µM) TCR:pMHC complexes with non-binding (≥ 100 µM) complexes from c259 TCR peptide mutant library.

complex affinity. Comparing the binding score of binding complexes (< 100 µM) with non-binding complexes (≥ 100 µM), we see a significant increase in the binding scores, ascertained by a one-sided t-test that resulted in a p-value of $6.59 \times 10^{-13}$ considering a significance level of 0.05. These results further support AF3's ability to detect disrupting mutations and match closely with the conclusions drawn from the analysis of the 572G TCR binding coeliac antigens.

### References

[1] Pradyot Dash et al. "Quantifiable Predictive Features Define Epitope Specific T Cell Receptor Repertoires". In: *Nature* 547.7661 (July 6, 2017), pp. 89–93. ISSN: 0028-0836. DOI: 10.1038/nature22383. PMID: 28636592. URL: https://www.ncbi.nlm.nih.gov/pmc/articles/PMC5616171/ (visited on 10/21/2022).

[2] Jose Cabezas-Caballero et al. "Generation of T Cells with Reduced Off-Target Cross-Reactivities by Engineering Co-Signalling Receptors". In: *Nature Biomedical Engineering* (Jan. 2, 2026), pp. 1–12. ISSN: 2157-846X. DOI: 10.1038/s41551-025-01563-w. URL: https://www.nature.com/articles/s41551-025-01563-w (visited on 04/02/2026).

[3] Jonas Birkelund Nilsson et al. "NetMHCpan-4.2: Improved Prediction of CD8+ Epitopes by Use of Transfer Learning and Structural Features". In: *Frontiers in Immunology* 16 (Aug. 7, 2025). ISSN: 1664-3224. DOI: 10.3389/fimmu.2025.1616113. URL: https://www.frontiersin.org/journals/immunology/articles/10.3389/fimmu.2025.1616113/full (visited on 04/20/2026).
